# Precision Functional Neuroimaging Reveals Individually Specific Auditory Responses in Infants

**DOI:** 10.1101/2025.08.04.667740

**Authors:** Julia Moser, Alyssa K. Labonte, Thomas J. Madison, Lana Hantzsch, Han H. N. Pham, Kimberly B. Weldon, M. Catalina Camacho, Rebecca F. Schwarzlose, Sanju Koirala, Jacob T. Lundquist, Sooyeon Sung, Cristian Morales Carrasco, Robert J. M. Hermosillo, Steven M. Nelson, Jed T. Elison, Damien A. Fair, Chad M. Sylvester

## Abstract

Adaptively responding to salient stimuli in the environment is a fundamental feature of cognitive development in early life, which is enabled by the developing brain. Understanding individual variability in how the brain supports this fundamental process is essential for uncovering neurodevelopmental trajectories and potential neurodevelopmental risks. In the present study, we used a precision functional imaging approach to probe activation in response to salient auditory stimuli and its relation to brain functional networks in individual infants. A minimum of 60 minutes of fMRI BOLD data with an auditory oddball paradigm were collected in ten infants with a mean postmenstrual age of 48 weeks. Results demonstrate the feasibility of performing a precision functional imaging study to investigate individual specific responses to salient stimuli in infants. While responses to the auditory oddball were consistent between individuals in auditory processing areas, responses across the rest of the brain differed across individuals in their magnitude and shape. Individual specific response patterns appeared to be relatively stable and differed from other participant’s response patterns, despite fluctuations across runs. Commonalities and differences between individuals demonstrated in this sample contribute to our understanding of how the developing brain instantiates processing of salient stimuli. Our findings suggest that during early development, early unimodal processing is well conserved across individuals, however subsequent perceptual processing is still being personally defined. In this context, individual specific response patterns could be a promising target for biomarkers of normative brain and cognitive development.

## Introduction

Human neonates possess remarkable cognitive skills, such as recognition of their native language (Mehler et al., 1988) or their mother’s voice (DeCasper & Fifer, 1980). Even before birth, in the last trimester of pregnancy, fetuses can detect novel sounds (Draganova et al., 2007) as well as learn basic auditory patterns (Moser et al., 2021; Schleger et al., 2014). This capacity of the developing brain to respond to salient stimuli in the environment is a fundamental aspect of cognitive development. Salient stimuli are stimuli that are either novel, unexpected, physically salient (e.g. loud or bright) or evolutionary significant and typically elicit attentional resources and prioritized processing. Characteristics of event-related responses towards novel stimuli in infancy can be associated with later cognitive outcomes (Katus et al., 2020; Weber et al., 2016), but also with risk factors such as high maternal anxiety (Sylvester et al., 2021). In this context, detecting differences in processing salient stimuli might be a key component towards characterizing variability in early brain functioning. Alterations in brain functioning in infancy are associated with children’s cognitive outcomes (Graham et al., 2016; Spann et al., 2018) but also with their risk for psychiatric disorders (Rogers et al., 2017; Sylvester et al., 2018). Yet, our knowledge about how the developing brain supports processing of salient stimuli is still limited. At the same time, understanding individual differences in this fundamental process might be essential for elucidating neurodevelopmental trajectories and potential risks for maladaptive outcomes.

Using functional Magnetic Resonance Imaging (fMRI), basic sensory tasks using motor (Dall’Orso et al., 2018) or auditory (Dehaene-Lambertz et al., 2002; Kosakowski et al., 2023; Sylvester et al., 2021) stimuli can be performed in sleeping infants. More advanced learning paradigms can be studied in older infants and toddlers during wakefulness (Ellis et al., 2021; Yates et al., 2025). These tasks can provide insights into the nature of information processing in the developing brain. For example Sylvester et al., (2021) found that auditory oddball stimuli elicit increased activity in many brain areas in newborns, including areas putatively belonging to the salience, cingulo-opercular, and ventral attention networks.

Characterizing the functional properties of brain regions that respond to salient stimuli in infants is thereby an essential step to understand the computations that are performed on incoming stimuli and even more importantly, how this processing changes with development. In adults, distributed sets of brain regions that perform related functions tend to show correlations in activity at rest (i.e. high ‘functional connectivity’). Such brain ‘functional networks’ have been well defined in adults and can be detected in the perinatal period (e.g. Sylvester et al., 2022). Functional networks show a developmental trend from late gestation (fetuses or preterm neonates) into infancy and childhood (Doria et al., 2010; Gao et al., 2015; Hu et al., 2022; Thomason et al., 2015) moving from a local to a distributed organization to increase processing efficiency (Fair et al., 2007).

The exact dimensions (size, boundaries, islands) of such functional connectivity networks and therefore the functionally relevant correlations in activity at rest are individual specific (Gordon et al., 2017; Laumann et al., 2015; Marek & Greene, 2021) and their patterns can even give insights about mental health (Cui et al., 2020; Demeter & Greene, 2024; Gratton et al., 2020; Labonte et al., 2024; Lynch et al., 2024). Recent research demonstrates this individual specificity in resting state networks as early as in newborn infants (Labonte et al., 2025; Molloy & Saygin, 2022; Moore et al., 2024; Myers et al., 2024). During this early phase of brain development, it is however unknown how these individual specific functional areas that co-fluctuate during rest are recruited during processing of incoming stimuli. Questions regarding the individual architecture of such networks and whether different infants recruit different networks during processing of the same stimulus remain less understood. At the same time, uncovering individual patterns of network activation during stimulus processing might be a key factor for understanding the associations between individual differences in event-related responses towards novel stimuli in infancy and neurodevelopmental outcomes.

Precision functional imaging is a strategy used to uncover individual patterns of functional brain organization. To study individual differences in how salient stimuli are processed by the infant brain, precision functional imaging can also be used to obtain personalized task activation maps. Individual specific mapping of task activation onto personalized networks with precision functional imaging is feasible in adults as demonstrated by Gordon et al., (2017), who showed that some of the correspondence between task-evoked activity and network organization might be missed without considering individual variation. Precision imaging requires large amounts of data for each individual, to maximize signal to noise and ensure reliability of individual specific activation patterns. Collecting large amounts of data is challenging in infants as the length of scanning sessions is limited by an infant’s comfort in the scanner without moving (Dubois et al., 2021; Korom et al., 2021; Labonte et al., 2024). In addition to refined practical methodological strategies for data acquisition, such as scheduling back-to-back appointments with families and working around infants’ bedtime routines, feasibility of precision functional imaging in infants can be increased by adopting advanced acquisition and processing strategies. For example, multi-echo (ME) data acquisition and additional thermal noise removal with NORDIC help to increase data reliability with fewer minutes of data compared to more traditional approaches (Moser et al., 2024). NORDIC denoising also increases functional contrast in task-based paradigms (Vizioli et al., 2021).

In the present study we use a precision functional imaging approach to probe activation in response to salient auditory stimuli and its relation to brain functional networks in individual infants. To achieve this, we assess task-based blood oxygen level dependent (BOLD) responses towards an auditory oddball, marking an unexpected, salient stimulus, in sleeping infants across multiple scanning sessions, building up on a group level study by Sylvester et al., (2021). We show the feasibility of this approach in an infant sample and additionally investigate the stability of task-based responses in individual infants across runs and acquisition days to determine the reliability of these responses. This work provides the foundation for studying individual differences in how the developing brain supports processing of salient stimuli and will help our understanding of individual differences in this fundamental process.

## Methods

### Sample

The sample for this study consists of ten infants enrolled at the University of Minnesota (mean postmenstrual age at first scan: 48 weeks (range: 45-51); 6 female). Mean gestational age at birth was 38.6 weeks (range 34-41) with three infants being born late-preterm. One of these had a hypoxic-ischemic encephalopathy at birth with no acute concerns during study participation. One study participant had a pregnancy-related indentation in the skull without concerns regarding the brain. Mean birthweight was 3352 g (range from 2041 g to 4141 g). Data included in this manuscript were collected in the framework of a study approved by the University of Minnesota’s Institutional Review Board, and written informed consent was obtained from both parents of the participants. Data for each participant were acquired over two to four days within one week. All data were acquired during natural sleep (Table 1).

**Table 1:** Overview of available data per participant. * initially acquired data for PB0019: 19 oddball runs (127min); 7 runs were excluded due to a hardware related artifact and one due to excessive motion. ** excludes one entire high motion run for PB0007 and two for PB0009, which are included in the task analysis with FD<0.9.

|  | Acquisition days | Oddball runs | Total time | Low motion data FD<0.9mm | Low motion data FD<0.3mm |
| --- | --- | --- | --- | --- | --- |
| PB0007 | 4 | 24 | 161 min | 153 min | 138 min** |
| PB0009 | 4 | 23 | 152 min | 142 min | 127 min** |
| PB0010 | 2 | 13 | 87 min | 84 min | 79 min |
| PB0011 | 3 | 11 | 74 min | 71 min | 67 min |
| PB0015 | 3 | 20 | 134 min | 126 min | 119 min |
| PB0016 | 2 | 12 | 80 min | 78 min | 75 min |
| PB0017 | 2 | 13 | 87 min | 84 min | 79 min |
| PB0018 | 3 | 15 | 98 min | 95 min | 92 min |
| PB0019* | 3 | 11 | 74 min | 67 min | 60 min |
| PB0020 | 2 | 13 | 87 min | 81 min | 76 min |

### Data Acquisition

Data was acquired with a 3T Siemens Prisma scanner with a 32 channel head coil. fMRI data were acquired with a four-echo version of the CMRR multiband (MB)-ME sequence (Feinberg et al., 2010; Moeller et al., 2010; TE = 14ms, 39ms, 64ms, 88ms, TR = 1.761s, 2mm resolution, MB factor = 6, flip angle = 68°). T2w and T1w anatomical references were also acquired (T2: TR = 42.3s, TE = 323ms, resolution = 0.8 × 0.8 × 0.8 mm, flip angle = 120°; T1: TR = 2.4s, TE = 2.2ms, resolution = 0.8 × 0.8 × 0.8 mm, flip angle = 8°) along with spin echo fieldmaps in both AP and PA direction for distortion correction (2 frames, TR = 8.4 s, TE = 66 ms, flip angle = 90°).

Infants were outfitted with multiple layers of hearing protection consisting of moldable silicone ear plugs, held in place by skin sensitive tape and OptoAcoustic headphones (OptoAcoustics Ltd., Tel Aviv) that reduced the sound of the scanner, while allowing us to provide an auditory stimulus. Each BOLD run lasted for 6.7 minutes and contained a passive listening auditory oddball paradigm. The oddball consisted of a 400ms long white noise stimulus, played in irregular intervals (10.5-14 seconds), marking a novel, deviant sound with respect to the regular scanner background noise. Each run started and ended with ~1 minute of regular scanner noise with 24 oddball stimuli in the middle (see also Sylvester et al., 2021). While trying to acquire as many BOLD runs as possible, scan time within one session was limited to a maximum of 60 minutes.

### Data Processing

Data were converted to BIDS format with dcm2bids (Boré et al., 2023). Each echo of the ME echo data was denoised with NORDIC (Dowdle et al., 2023; Moeller et al., 2021; Vizioli et al., 2021). Phase and magnitude images of the scan were used for NORDIC as well as three noise frames acquired at the end of each functional run to help estimate the empirical thermal noise level. NORDIC was implemented in Matlab R2019a. Brain tissue segmentations of the anatomical data were generated using BIBSNet (Hendrickson et al., 2024), a deep learning model trained on hand edited segmentations of 0-8 month old infants. All data were preprocessed using NiBabies 25.0.1 (Goncalves et al., 2025). The options “--multi-step-reg” and “--norm-csf” were used to facilitate registration of native space anatomical data to standard space and M-CRIB-S (Adamson et al., 2020) was used for surface reconstruction (“--surface-recon-method mcribs”). For functional data processing the option “--project-goodvoxels” was enabled to exclude voxels with locally high coefficient of variation from volume-to-surface projection of the BOLD time series, and “--cifti-output 91k” to enable output of BOLD time series and morphometric data (surface curvature, sulcal depth, and cortical thickness maps) in the HCP grayordinates space (Glasser et al., 2013). Susceptibility distortion correction of BOLD time series was done within NiBabies, using SDCFlows with an FSL topup-based method to estimate fieldmaps from “PEPolar” acquisitions (acquisitions with opposite phase encoding direction; Andersson et al., 2003). For four runs of PB007 no fieldmaps were available matching the BOLD data based on problems during data acquisition. For these runs, the experimental “test-syn” version of NiBabies was used with the “--use-syn-sdc” flag, which uses a synthetic fieldmap correction approach. Data post-processing was done separately for task analysis and functional connectivity analysis. In both instances, postprocessing was performed using XCP-D 0.10.5 (Mehta et al., 2023) with the “hbcd” mode for setting defaults and the “--motion-filter-type” set to “none” as the TR of the scanning sequences used here were too slow to adequately resolve infants’ fast breathing rate (30-60bpm). Options used for task analysis were “--lower-bpf 0.025”, “--upper-bpf 0.15”, “-FD=0.9” and “--nuisance-regressors none” as nuisance regressors were instead included in the general linear model (GLM) for task analysis. Options used for the connectivity analysis were “--lower-bpf 0.009” (instead of the default 0.01), the default “--upper-bpf” of 0.08, the default nuisance regressors and “-FD=0.3”. Runs with less than 60% low motion data were excluded from further analysis as well as runs with hardware related artifacts or excessive motion leading to unusable alignment during preprocessing. XCP-D outputs used for task analysis were furthermore checked for grayordinates containing NaN’s, which were then spatially interpolated and data were smoothed with a smoothing kernel of 2.25 using workbench command “-cifti-smoothing”.

## Data Analysis

### Task analysis

Infant task responses were mapped using a GLM implemented in Nilearn (Nilearn contributors et al., 2025). The first level model included the oddball as a main effect. A standard double gamma canonical hemodynamic response function (HRF; “glover”) was used to model task activation. The function’s derivative was additionally included in the model to add flexibility (Friston et al., 1998) as HRFs are known to vary in infants (Arichi et al., 2012). Frames that exceeded the FD threshold of 0.9 mm (Siegel et al., 2014) were excluded from the analysis (scrubbed from the data and design matrix). Significance in each participant was determined by calculating a noise distribution. This noise distribution contained 1000 permutations of the same analysis with a randomly scrambled design matrix instead of the design matrix matching the events. The significance threshold of p<0.01 was equal to the 0.5th and 99.5th percentile of this distribution. For the exploratory model-free analysis we used the same first level model in Nilearn with a Finite Impulse Response (FIR) Function, modeling frames 1-11 after oddball onset. In an additional analysis, the first 8, middle 8 and last 8 oddballs were coded and modeled individually to investigate effects of habituation within a run (Sylvester et al., 2021). Suppl. Figure 1 contains an example for all three design matrices.

BOLD time courses which were used for inspection of event-related responses were created from the concatenated time series outputted by XCP-D. Time series were averaged across a region of interest and then this signal was averaged across all event onsets. Furthermore, stability of oddball responses across runs within an individual were determined. Split halves were created for each participant by comparing responses to half their runs (leaving one out for an odd number of runs) to the other half. All unique run combinations were computed 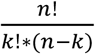 with n being the number of available runs and k the number of runs used in one half with 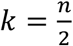 rounded to the smaller integer) and the least number of possible combinations (462 for n=11) used as the number of permutations per participant. The dice coefficient was used as a metric of overlap between split halves for each permutation, using the dice overlap of the 25% highest beta values. Overlaps across permutations were counted with again overlapping these overlaps.

### Network analysis

Individual specific networks were created using template matching (Hermosillo et al., 2024) with the infant specific approach used by Moore et al., (2024). To mitigate the impact of the task on the organization of these functional networks, only data before the first and after the last oddball from each run was used. Furthermore, data with FD>0.3mm were excluded for creation of the functional connectivity matrices underlying these analyses. To calculate the percent of a network being active during the task, vertices assigned to a network were counted and their fraction of active vertices calculated. In this analysis active vertices were defined as the highest 25% of beta values. The infant specific template published by Moore et al., (2024) is restricted to surface vertices, which also restricts this analysis to cortical grayordinates.

## Results

### Precision imaging data reveals individual specific oddball responses

We acquired a minimum of 60 minutes of low motion data from each of the ten study participants (Table 1). Total acquired data ranged from 74-161 min with a mean of 103 min, excluding 8 oddball runs from PB0019. Data included in the task analysis (FD<0.9) ranged from 67-153 min (mean = 98 min; Table 1; Suppl Fig 2). Data included in the connectivity analysis (FD<0.3) ranged from 60-138 min (mean = 91 min; Table 1; Suppl Fig 3), excluding two entire runs for PB009 and one for PB007 due to not meeting the 60% low motion requirement.

Each participant showed a significant response to the oddball stimuli (p<0.01) using data from all acquired runs. Thereby response patterns were individual specific for each participant (Figure 1). Z-scored beta values across the whole brain are displayed in Suppl. Figure 4. The fraction of brain areas being involved in processing the oddball stimulus varied greatly between participants, from 5% of grayordinates showing positive significant activation to 65%. Those participants with larger portions of the brain being active while processing the oddball stimulus showed higher overall beta magnitudes (Figure 1; Suppl. Figure 4). Even when strictly selecting areas of largest activity (75th percentile of beta values) - to get another estimate independent of significance thresholds - individual specificity remained.

**Figure 1:**
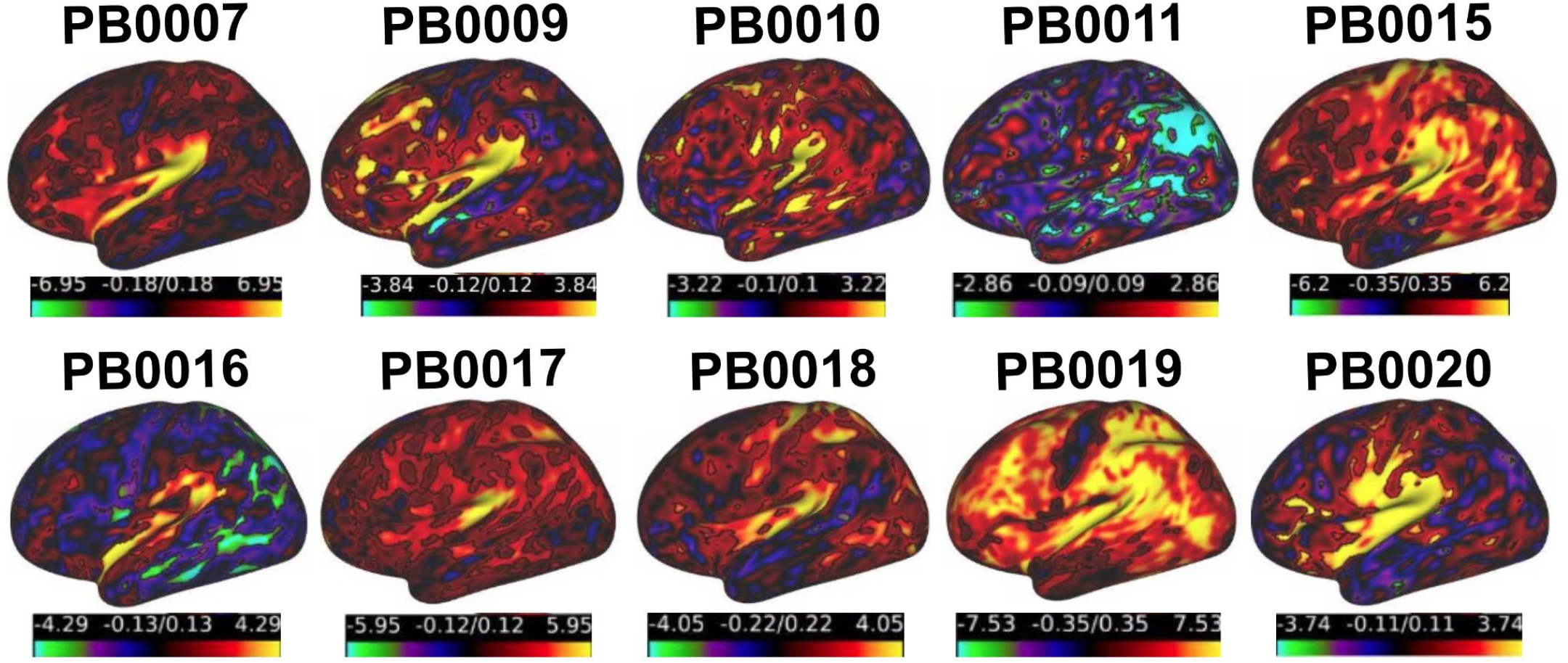
Z-scored beta values showing individual patterns of activation towards the oddball stimuli. Significant areas are outlined in black. Values are scaled from 5th to 95th percentile for each individual signal range. Full maps are displayed in Suppl. Figure 4.

In the context of precision imaging, the question arises whether the individual specific response patterns we see in these participants reflect meaningful individual traits or whether they are a product of variability between acquired runs and are more state-like features or even dominated by other factors in the data such as motion. To investigate stability across runs, we conducted a permutation analysis within subjects, comparing overlap between beta maps of all possible unique data split halves (by run). On average, the overlap between the top 25% beta values in split halves of the data was moderate (overall mean = 0.32, ranging from a mean of 0.5 in PB007 to 0.12 in PB0011; Figure 2). Despite the moderate overlap between split halves, for seven out of ten participants, the overlap with the second half of their own dataset was significantly higher compared to the overlap with the data halves of other participants (p<0.001 ‘self’ vs. ‘other’; Figure 2). PB010 and PB018 showed non-significant differences, while PB011 had significantly less overlap with its own second halves. As PB0011 showed the largest amounts of significant negative activation, we repeated the analysis for PB0011 with the 25% lowest negative betas, in which case overlap with the second half of their own dataset was significantly higher compared to the overlap with the data halves of other participants (p<0.001; Suppl. Figure 5A). Furthermore, all subjects showed distinct brain areas that showed overlap between split halves in close to 100% of permutations (Suppl. Figure 6 and 5B). This underlines the existence of individual specific response patterns in the majority of cases.

**Figure 2:**
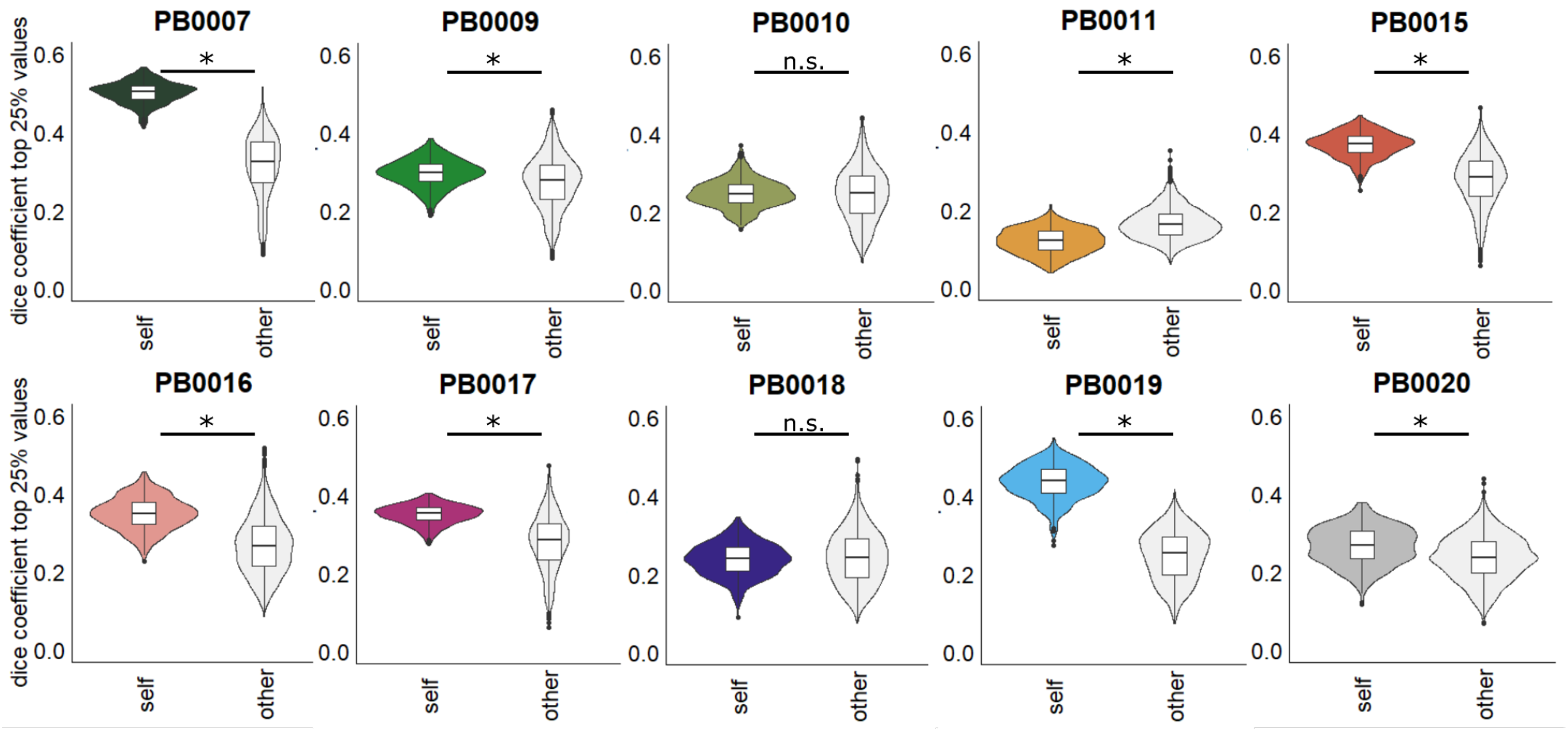
Dice overlap of top 25% beta values calculated from split halves in 462 unique combinations of runs for each subject. Comparisons are between split halves (colored) and the first half of every other subject (light gray). * denotes significance with p<0.001.

Differences in stability between participants did not show an obvious relationship to the number of runs included or a participant’s motion metrics (Suppl. Figure 7A-C). Comparing differences in stability for various run combinations within participants, some participants showed lower stability for run combinations in which higher motion runs were unequally distributed between split halves while for others this increased stability or had no effect (Suppl. Figure 7D). This speaks for a complex, not apparently systematic relationship between motion and task activation stability in this sample. High motion runs also did not systematically contribute more or less to subjects’ individual specific activity patterns (Suppl. Figure 8).

The variability in magnitudes of beta values between participants remained when using an unassumed response to model BOLD activation after the oddball onset. Therefore it can not fully be accounted for by a subject specific fit to the assumed response used in this analysis but rather represents an individual specific degree of activation towards the oddball (Suppl. Figure 9A). Furthermore, the number of removed frames right after the onset of an oddball, which impact task modeling, were overall low and differences between participants not associated with beta magnitudes (Suppl. Figure 9B).

### Individual responses overlap in key areas while recruiting different network patterns

Across individuals in our sample, oddball responses overlapped in core auditory processing areas such as the left and right auditory cortices, the thalamus and inferior colliculus as well as parts of visual cortex and the supplementary motor area (Figure 3). Figure 3 shows the overlap in significant responses from the first level model across all ten participants. Suppl. Figure 10 demonstrates that the pattern is very similar when ignoring significance but looking at the top 25% of values in each subject instead. Task-related responses extracted from subject’s fMRI time courses and time-locked to oddball onsets demonstrate synchronicity across participants in areas of high and medium overlap in the first level model (Figure 4). In areas of low overlap, participants varied in the timing and amplitude of the stimulus related BOLD signal change, which was non-significant in some participants and even negatively associated with the assumed response in others (see PB0011 in Figure 1 and Suppl. Figure 4). Variance in timing of the first peak after the oddball was significantly lower in the example regions with high overlap (var = 0.168) compared to regions with medium (var = 0.523) or low overlap (var = 0.503; F(2,7) = 5.19; p=0.02). These analyses show high overlap between individuals for brain regions primarily associated with auditory processing while demonstrating individual variability further along the processing hierarchy. In the within subject permutation analysis, many of the areas that are highly stable within a participant, form a similar pattern as the overlap across participants, with primary auditory areas and some visual areas showing the greatest degree of stability (Suppl. Figure 6).

**Figure 3:**
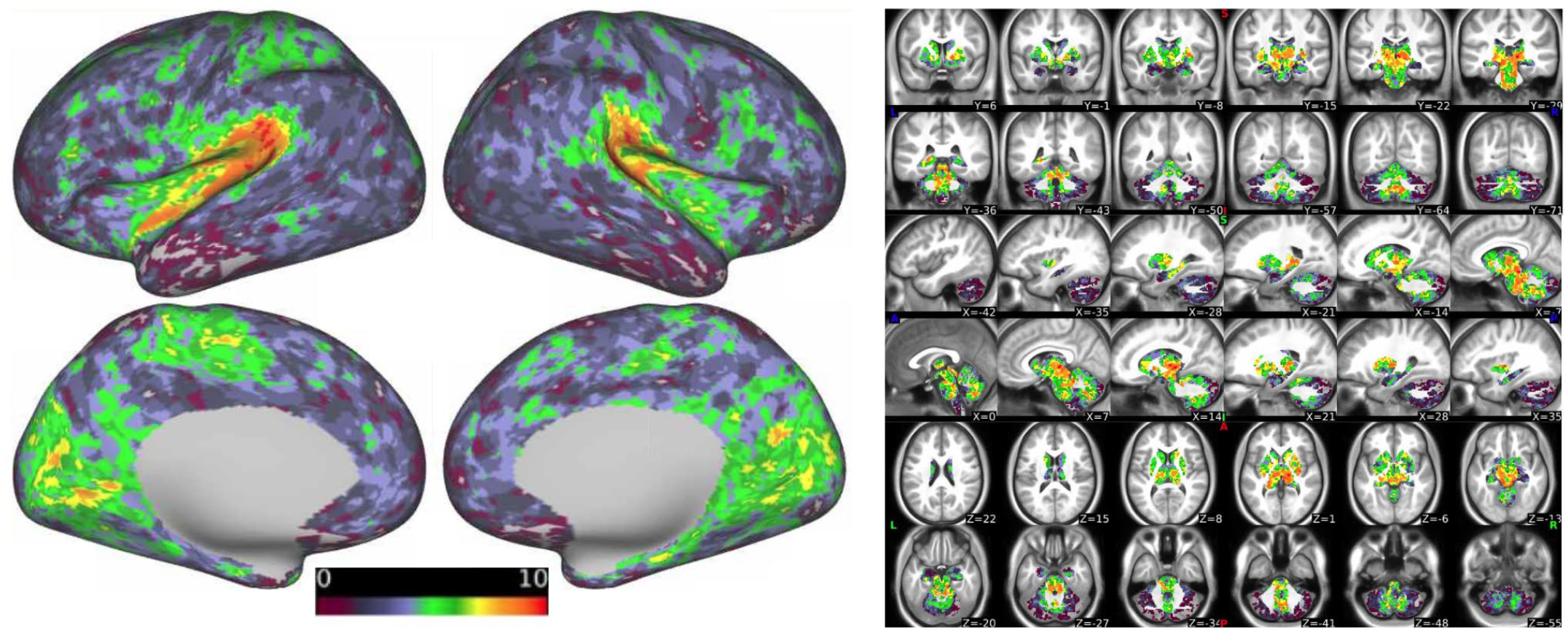
Overlap of areas with significant activation (p<0.01) across individuals. Scale 0-10 is the participant count. Overlap is the highest in core auditory processing areas.

**Figure 4:**
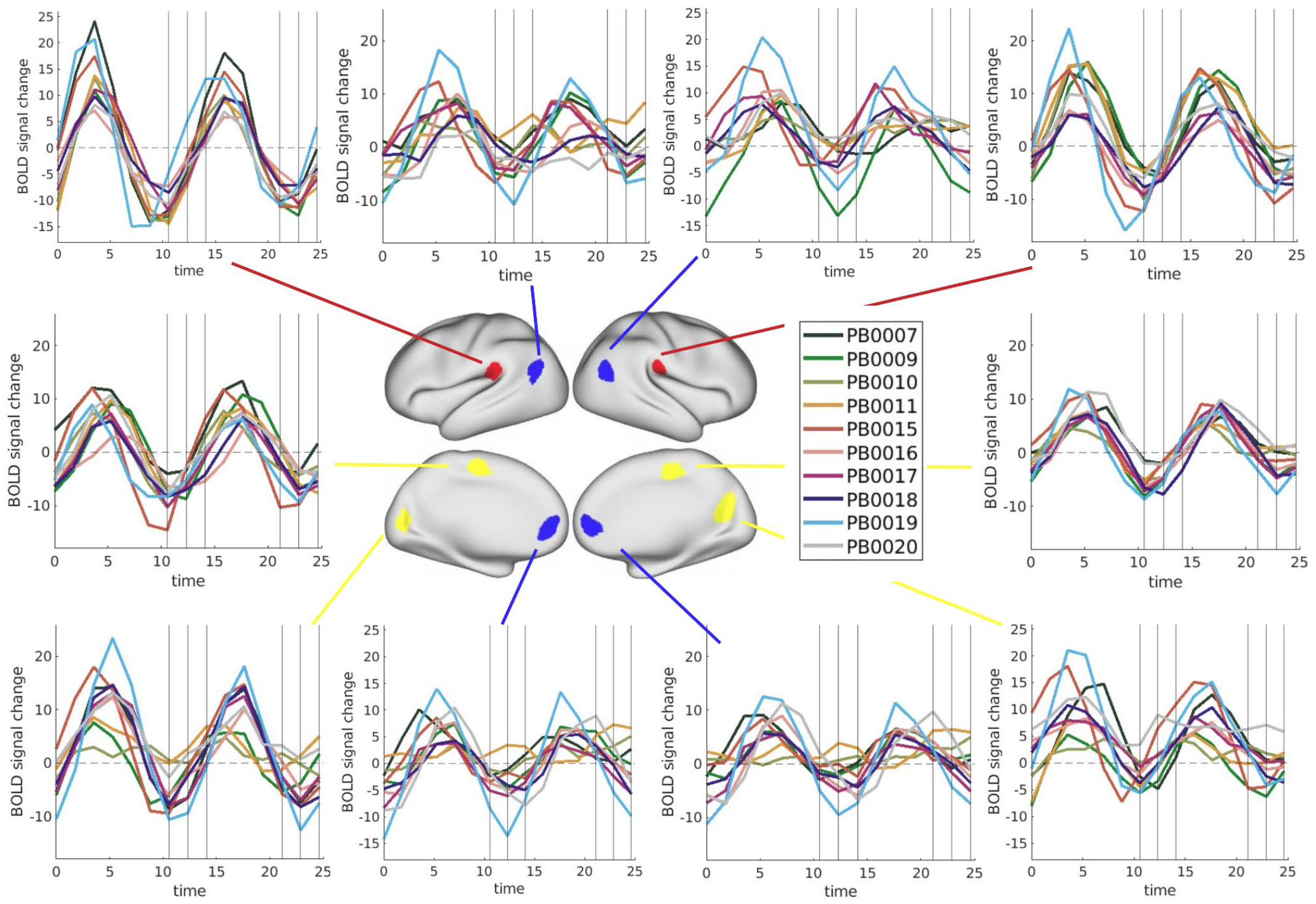
Individual time courses (trial averaged BOLD signal) for regions of interest with high (red), middle (yellow) and low (blue) overlap (see Figure 3). Vertical lines indicate the onset of the following oddball (jittered by 1-3 TRs).

Overlapping each individual’s areas of strongest activation towards the oddball (75th percentile) with their individual specific functional connectivity networks, the auditory network showed the highest percentage of activation in every case (Figure 5 & Suppl. Figure 11). The second largest percentage of activation could be seen in the parietal-occipital network (PON) in half the participants while the other half did not share a consistent second largest network. Each participant showed an individual specific pattern of networks that were recruited to process the oddball stimuli (Suppl. Fig 11). Restricting the network analysis to the areas that show high stability across run combinations (areas that overlap between split halves and between more than 50% of run combinations), the auditory network was still the network with the strongest involvement across all participants but two (PB0018, PB0011) for whom the visual network dominated. Despite stable areas being smaller than the areas used in the initial analysis, the individual specific pattern of networks that were recruited within each participant remained in all participants but PB0011 who showed very little remaining stable activation (Suppl. Figure 12 & 13).

**Figure 5:**
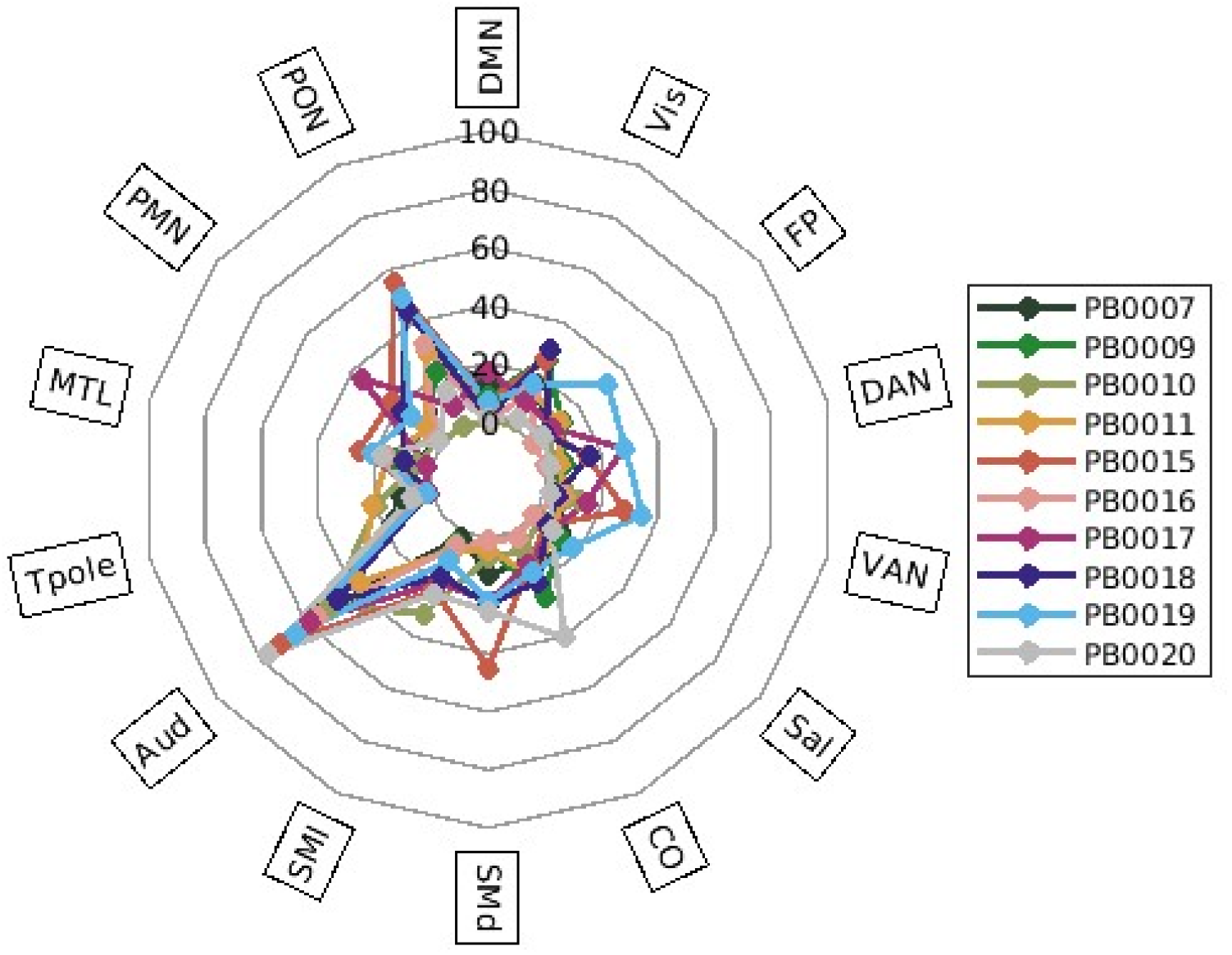
Percent of a network containing high beta values (top 25%) for each participant. DMN: default mode network; Vis: visual network; FP: fronto-parietal network; DAN: dorsal attention network; VAN: ventral attention network; Sal: salience network; CO: cingulo-opercular network; SMd: somato-motor dorsal; SMl: somato-motor lateral; Aud: auditory network; Tpole: temporal pole; MTL: medial temporal lobe; PMN: parietal memory network; PON: parietal-occipital network.

Based on prior results (Sylvester et al., 2021) we expect that BOLD responses towards the oddball stimuli to change/become attenuated across the 24 repetitions within one run. This might account for some of the variability in responses as the nature of this change might be different between participants. In a follow up analysis, we compared the degree of overlap between participants when only coding for the early (first 8) or late (last 8) oddballs in a run. Separating out responses towards only first and only last oddballs decreased the overlap between participants compared to using all events. Using all events, there was high overlap between participants in 31% of vertices (quantified as active in 50% of the sample or more; Figure 3) which reduced to 28% using only early oddballs and to 4% with only late oddballs (Suppl. Figure 14). Responses towards the late oddballs overall declined, with less areas with significant activation (mean percent of active grayordinates across subjects for early = 34.36 (SD = 19.46) and late = 17.72 (SD = 10.73)), replicating the habituation effect found in Sylvester et al., (2021).

## Discussion

This study demonstrated the feasibility of performing a precision functional imaging study to investigate individual specific responses towards salient stimuli in infants. While responses to the auditory oddball could be detected in each individual infant, they differed in their magnitude and shape across the brain. Areas with the most overlap across individuals were those belonging to the auditory network, followed by parietal-occipital areas. In these areas with high overlap, average time courses were highly synchronous across individuals. Using subsets of the data in a within-subject permutation test, we could show that individual specific response patterns were relatively stable and differed from other participants’ response patterns in the majority of cases. Areas that were most stable within individuals were similar to areas with the most overlap across individuals.

### Auditory task mapping in individuals

With the auditory oddball paradigm used in the present study, we were able to map an individual’s brain responses to salient stimuli. In each individual, auditory oddball responses mapped onto auditory areas including the primary auditory cortex and the anterior insula, a region well characterized for processing multimodal salient stimuli. Highest overlap between participants in areas associated with auditory processing (inferior colliculus, thalamus, primary auditory cortex) underline the importance of these primary auditory areas in novelty and salience detection. This is in line with studies investigating the processing hierarchy of auditory prediction errors in human adults (Wacongne et al., 2011) and non-human primates (Chao et al., 2018; Jiang et al., 2022; Uhrig et al., 2014). Outside of auditory regions, the parietal-occipital network (PON), a network along the dorsal visual stream, showed the most consistency across participants. Even though this appears surprising at first glance for an auditory paradigm, it is in line with meta-analytic findings in adults showing consistent involvement of part of the medial visual cortex across auditory oddball studies (Kim, 2014), an area broadly involved in action planning and control and spatial navigation (Hutchison et al., 2015). In this infant sample, a still increased cross-modal innervation of fiber tracts could be an additional contributor to this pattern (von Melchner et al., 2000).

While the auditory network had the largest percent of activation in all subjects, the rest of the response pattern was individual specific, even across different combinations of runs. For example, in PB0017 larger fractions of the parietal memory network and dorsal attention network were active compared to other networks, while for PB0009 and PB0020 the cingulo-opercular network played a more important role. As mentioned above, the PON played an important role for many subjects but activation in auditory and PON was for example combined with somato-motor dorsal in PB0015, visual in PB0018 and fronto-parietal and ventral attention in PB0019. This variety in network patterns across individuals hints towards differences between individual infants in terms of the areas that they recruit to process a novel, salient stimulus. One underlying reason for this could be a refinement in network function that is still taking place, where some roles are not yet clearly distributed and pathways will get further shaped with prolonged exposure to the environment. At the age this sample was measured, systems are still highly plastic and can take over different roles. For example, a study in congenitally blind individuals showed an important role of the visual system in processing non-visual information (Leo et al., 2012) similar to animal studies demonstrating cross-modal innervation when appropriate extrinsic input is missing (Laemle et al., 2006; von Melchner et al., 2000).

While this is purely speculative for now, future longitudinal studies can help to investigate if this variety in task activation patterns is a phenomenon just occurring during this early, highly plastic phase of brain development and patterns get more similar across time or whether these patterns stay with individuals. If response patterns get more similar across time, this would highlight the important role of extrinsic inputs in shaping the brain’s functional organization, particularly in individuals exposed to similar environments. If patterns are staying with individuals, a person’s specific response patterns towards a salient stimulus, could be a promising target for a biomarker of normative brain and cognitive development. If response patterns toward salient stimuli in an infant’s environment already take different shapes in early infancy, this will impact the cascade of cognitive development and might be one potential mechanism for how differences in brain functional architecture relate to behavioral outcomes. Similar to how under or over responding to a salient stimulus might be maladaptive, recruitment of a specific network pattern might be particularly adaptive during a distinct developmental phase. Understanding individual differences in this fundamental process could be an essential step for uncovering neurodevelopmental trajectories and potential risks in terms of mental health outcomes.

### Considerations for infant precision neuroimaging with task paradigms

When using a task design in the context of an individual specific precision imaging study, stability of the metric under investigation plays a crucial role. In our within-subject permutation analysis, we found overlap between split halves to be moderately high across different run combinations. Looking into the areas that show overlap within and across permutations (Suppl. Figure 6), a relatively large degree of within-subject variability remains, mainly outside of areas primarily associated with auditory processing. Despite this variability, patterns remain individual specific. This speaks towards some ‘core’ individual processing strategies that can be identified despite the variability added by noise, motion or other physiological factors. Biological factors such as sleep stage can for example impact responses to the oddball. Prior studies using auditory stimuli highlighted the differences in stimulus processing between active and quiet sleep (Dehaene-Lambertz et al., 2002; Moser et al., 2020), which is a variable not tracked in this dataset. Association of active sleep with increased motility (Thoman & Tynan, 1979) furthermore complicates disentangling effects of behavioral states from motion artifacts.

This within-subject variability that is likely caused by a mix of biological factors and noise is important to keep in mind when evaluating task activation in a precision imaging context. When evaluating resting state functional connectivity, more data is usually helpful to most accurately represent individual patterns and a plateau in reliability is reached given enough time. Our results in this task analysis context did not indicate that more runs necessarily lead towards more consistency (e.g. comparing dice overlap results from PB0009 with 23 runs and PB0019 with 11 runs) and it is necessary to further explore factors causing variability between runs such as the behavioral state differences discussed above. The precision imaging style data acquisition across multiple days and often with multiple attempts within one session could for example induce variability based on placement in the head coil (King et al., 2023). Another potentially relevant variable could be timing and mode of feeding before the scan (Liu et al., 2000; Zhang et al., 2022). Based on our observations, we suggest that consistency analysis within subjects should be implemented as a part of a precision imaging task paradigms.

Another important consideration for using task-based BOLD activation in a precision imaging context is the variability or BOLD hemodynamic responses themselves. While BOLD responses are stable within an individual across prolonged scan times or sessions, they vary across individuals and brain regions (Aguirre et al., 1998; Menz et al., 2006) in addition to variations by age (Arichi et al., 2012; Rieck et al., 2022). Along with contributions by noise or state dependent physiological factors, these individual specific variations in BOLD responses potentially contributed to the differences in magnitude and extent of activation we observed in our sample. This complicates finding individual specific thresholds for meaningful activation towards a stimulus. In the present study, we tried to circumvent this by focusing on areas with the highest activation (in percent). This allowed us to address questions around the spatial patterns of oddball responses independent of individual response magnitudes, which showed consistency across modeling with an assumed response or an FIR model.

### Limitations and Outlook

This is the first study of its kind, demonstrating feasibility of precision functional imaging with a task in infants, and at the same time facing several limitations. Our sample was small and not homogeneous, both in terms of postmenstrual age and gestational age at birth, which precludes interpretation of any differences between subjects. While our individual specific analyses still contribute to our understanding of task activation patterns in infants, future studies should try to look at trajectories within individuals and leverage longitudinal designs to untangle between subject differences and developmental effects.

Measuring fMRI BOLD activation in a task paradigm is furthermore always limited by our accuracy of modeling hemodynamic responses. The method used in the present paper is a standard approach, which is not specifically tailored to infant studies. While inclusion of the HRF’s derivative contributes to capturing differences in the response’s shape, it is unlikely that this approach provides the optimal fit for every infant. Work in newborns provided evidence that neurovascular coupling may not even be fully developed (Hendrikx et al., 2019), which further complicates our assumptions about hemodynamic responses in our infant sample. Using an unassumed response with an FIR model is however more difficult for individual based analyses in precision imaging as the traditional approach to determine significant activation by measuring an effect of time across all modeled frames is only well powered at a group level. Fitting more flexible HRF functions which are specific for individuals and different parts of the brain like for example with an HRF library (Prince et al., 2022) will be an important future step for task precision imaging in developmental populations. During this early phase of exploring analyses of individual specific task responses as markers of stimulus processing capabilities in early development, it could furthermore be helpful to consider alternative task designs. While for example a block design approach is not feasible for the question under investigation here - processing of a salient stimulus - it might be helpful for other research questions around sensory processing as it increases stability and facilitates modeling.

For mapping task activation to individual networks, we used an infant specific version of template matching (Moore et al., 2024). While the template used here is specific to an infant sample, it was generated from a slightly younger age group using a different data acquisition sequence. Future work could explore further options to get more independent individual specific network solutions. Only segments of the data which did not contain events were used for generating functional connectivity networks to resemble resting state as well as possible. While tasks do not have a major impact on individual specific network patterns (Gratton et al., 2018), they change the frequency content of BOLD signals (Bailes et al., 2023), which could have had an impact on our ‘resting state’ data.

Lastly, to be better able to disentangle different contributors to fluctuations of activation patterns between different run combinations, future task-based precision imaging research should try to collect additional variables such as respiration or heart rate to have a better understanding of activity states and physiological differences between task runs.

## Conclusions

With demonstrating the feasibility of performing a precision functional imaging study to investigate individual specific responses towards salient stimuli in infants, this study opens the doors for individual specific task assessments in early development. In this auditory task, high synchronicity between participants in areas involved in auditory processing highlighted basic mechanisms that are common across individuals while variability in other active areas demonstrated individual specific recruitment of additional areas to process a novel, salient stimulus. Findings regarding stability across different run combinations provided evidence for fluctuations in task responses across runs which need to be accounted for when mapping individual specific task response patterns, for example by looking at within subject central tendencies. The commonalities and individual differences demonstrated in this sample contribute to shaping our understanding of how the developing brain enables processing of salient stimuli which is a fundamental skill during development. Results presented in this paper contribute towards the development of personalized biomarkers of normative brain and cognitive development.

## Data and Code Availability

Data can be made available upon request, given a formal data sharing agreement is set up by the institutions involved.

Code is available from the following repository: https://github.com/DCAN-Labs/infant_oddball_precision_imaging

## Author Contributions

Julia Moser: Conceptualization, Investigation, Methodology, Formal analysis, Visualization, Writing - Original Draft; Alyssa K. Labonte: Conceptualization, Writing - Review & Editing; Thomas J. Madison: Formal analysis, Software; Lana Hantzsch: Project administration, Data Curation; Han H. N. Pham: Data Curation; Kimberly B. Weldon: Software, Writing - Review & Editing; M. Catalina Camacho: Writing - Review & Editing; Rebecca F. Schwarzlose: Writing - Review & Editing; Sanju Koirala: Writing - Review & Editing; Jacob T. Lundquist: Software; Sooyeon Sung: Investigation, Writing - Review & Editing; Cristian Morales Carrasco: Methodology, Writing - Review & Editing; Robert J. M. Hermosillo: Conceptualization, Writing - Review & Editing; Steven M. Nelson: Writing - Review & Editing; Jed T. Elison: Resources, Writing - Review & Editing; Damien A. Fair: Conceptualization, Resources, Supervision, Writing - Review & Editing; Chad M. Sylvester: Conceptualization, Supervision, Writing - Review & Editing

## Funding

Individual author funding includes: DFG German Research Foundation 493345456 (author JM; Deutsche Forschungsgemeinschaft) and National Institute of Mental Health (R01MH134966 to JM, DAF, and CMS; R01MH122389 to CMS).

## Declaration of Competing Interests

Damien A. Fair is a patent holder on the Framewise Integrated Real-Time Motion Monitoring (FIRMM) software. He is also a co-founder of Turing Medical Inc that licenses this software. The nature of this financial interest and the design of the study have been reviewed by two committees at the University of Minnesota. They have put in place a plan to help ensure that this research study is not affected by the financial interest. Steven M. Nelson consults for Turing Medical, which commercializes FIRMM. This interest has been reviewed and managed by the University of Minnesota in accordance with its Conflict of Interest policies. The other authors declare no competing interests.

## Acknowledgements

We thank all the families of our precision babies for their participation and their dedication towards research. We also thank Bolade Santos, Isabella Linder, Angel Gathumbi and Katelyn Day for their support during data acquisition. The authors acknowledge the Minnesota Supercomputing Institute (MSI) at the University of Minnesota for providing resources that contributed to the research results reported within this paper. URL: http://www.msi.umn.edu

## References

Adamson, C. L., Alexander, B., Ball, G., Beare, R., Cheong, J. L. Y., Spittle, A. J., Doyle, L. W., Anderson, P. J., Seal, M. L., & Thompson, D. K. (2020). Parcellation of the neonatal cortex using Surface-based Melbourne Children’s Regional Infant Brain atlases (M-CRIB-S). Scientific Reports, 10(1), 4359.

Aguirre, G. K., Zarahn, E., & D’esposito, M. (1998). The variability of human, BOLD hemodynamic responses. NeuroImage, 8(4), 360–369.

Andersson, J. L. R., Skare, S., & Ashburner, J. (2003). How to correct susceptibility distortions in spin-echo echo-planar images: application to diffusion tensor imaging. NeuroImage, 20(2), 870–888.

Arichi, T., Fagiolo, G., Varela, M., Melendez-Calderon, A., Allievi, A., Merchant, N., Tusor, N., Counsell, S. J., Burdet, E., Beckmann, C. F., & Edwards, A. D. (2012). Development of BOLD signal hemodynamic responses in the human brain. NeuroImage, 63(2), 663–673.

Bailes, S. M., Gomez, D. E. P., Setzer, B., & Lewis, L. D. (2023). Resting-state fMRI signals contain spectral signatures of local hemodynamic response timing. eLife, 12, e86453.

Boré, A., Guay, S., Bedetti, C., Meisler, S., & GuenTher, N. (2023). Dcm2Bids (Version 3.1.1). 10.5281/zenodo.8436509

Chao, Z. C., Takaura, K., Wang, L., Fujii, N., & Dehaene, S. (2018). Large-scale cortical networks for hierarchical prediction and prediction error in the primate brain. Neuron, 100(5), 1252–1266.e3.

Cui, Z., Li, H., Xia, C. H., Larsen, B., Adebimpe, A., Baum, G. L., Cieslak, M., Gur, R. E., Gur, R. C., Moore, T. M., Oathes, D. J., Alexander-Bloch, A. F., Raznahan, A., Roalf, D. R., Shinohara, R. T., Wolf, D. H., Davatzikos, C., Bassett, D. S., Fair, D. A., … Satterthwaite, T. D. (2020). Individual Variation in Functional Topography of Association Networks in Youth. Neuron, 106(2), 340–353.e8.

Dall’Orso, S., Steinweg, J., Allievi, A. G., Edwards, A. D., Burdet, E., & Arichi, T. (2018). Somatotopic mapping of the developing sensorimotor cortex in the preterm human brain. Cerebral Cortex (New York, N.Y.: 1991), 28(7), 2507–2515.

DeCasper, A. J., & Fifer, W. P. (1980). Of human bonding: newborns prefer their mothers’ voices. Science (New York, N.Y.), 208(4448), 1174–1176.

Dehaene-Lambertz, G., Dehaene, S., & Hertz-Pannier, L. (2002). Functional neuroimaging of speech perception in infants. Science, 298(5600), 2013–2015.

Demeter, D. V., & Greene, D. J. (2024). The promise of precision functional mapping for neuroimaging in psychiatry. Neuropsychopharmacology: Official Publication of the American College of Neuropsychopharmacology. 10.1038/s41386-024-01941-z

Doria, V., Beckmann, C. F., Arichi, T., Merchant, N., Groppo, M., Turkheimer, F. E., Counsell, S. J., Murgasova, M., Aljabar, P., Nunes, R. G., Larkman, D. J., Rees, G., & Edwards, A. D. (2010). Emergence of resting state networks in the preterm human brain. Proceedings of the National Academy of Sciences of the United States of America, 107(46), 20015–20020.

Dowdle, L. T., Vizioli, L., Moeller, S., Akçakaya, M., Olman, C., Ghose, G., Yacoub, E., & Uğurbil, K. (2023). Evaluating increases in sensitivity from NORDIC for diverse fMRI acquisition strategies. NeuroImage, 270, 119949.

Draganova, R., Eswaran, H., Murphy, P., Lowery, C., & Preissl, H. (2007). Serial magnetoencephalographic study of fetal and newborn auditory discriminative evoked responses. Early Human Development, 83(3), 199–207.

Dubois, J., Alison, M., Counsell, S. J., Hertz-Pannier, L., Hüppi, P. S., & Benders, M. J. N. L. (2021). MRI of the Neonatal Brain: A Review of Methodological Challenges and Neuroscientific Advances. Journal of Magnetic Resonance Imaging: JMRI, 53(5), 1318–1343.

Ellis, C. T., Skalaban, L. J., Yates, T. S., & Turk-Browne, N. B. (2021). Attention recruits frontal cortex in human infants. Proceedings of the National Academy of Sciences of the United States of America, 118(12). 10.1073/pnas.2021474118

Fair, D. A., Dosenbach, N. U. F., Church, J. A., Cohen, A. L., Brahmbhatt, S., Miezin, F. M., Barch, D. M., Raichle, M. E., Petersen, S. E., & Schlaggar, B. L. (2007). Development of distinct control networks through segregation and integration. Proceedings of the National Academy of Sciences of the United States of America, 104(33), 13507–13512.

Feinberg, D. A., Moeller, S., Smith, S. M., Auerbach, E., Ramanna, S., Gunther, M., Glasser, M. F., Miller, K. L., Ugurbil, K., & Yacoub, E. (2010). Multiplexed echo planar imaging for subsecond whole brain FMRI and fast diffusion imaging. PloS One, 5(12), e15710.

Friston, K. J., Josephs, O., Rees, G., & Turner, R. (1998). Nonlinear event-related responses in fMRI. Magnetic Resonance in Medicine, 39(1), 41–52.

Gao, W., Alcauter, S., Elton, A., Hernandez-Castillo, C. R., Smith, J. K., Ramirez, J., & Lin, W. (2015). Functional Network Development During the First Year: Relative Sequence and Socioeconomic Correlations. Cerebral Cortex, 25(9), 2919–2928.

Glasser, M. F., Sotiropoulos, S. N., Wilson, J. A., Coalson, T. S., Fischl, B., Andersson, J. L., Xu, J., Jbabdi, S., Webster, M., Polimeni, J. R., Van Essen, D. C., Jenkinson, M., & WU-Minn HCP Consortium. (2013). The minimal preprocessing pipelines for the Human Connectome Project. NeuroImage, 80, 105–124.

Goncalves, M., Moser, J., Madison, T. J., McCollum, R., Lundquist, J. T., Fayzullobekova, B., Hadera, L., Pham, H. H. N., Moore, L. A., Houghton, A. M., Conan, G., Styner, M. A., Alexopoulos, D., Smyser, C. D., Stoyell, S. M., Koirala, S., Nelson, S. M., Weldon, K. B., Lee, E., … Fair, D. A. (2025). FMRIPrep Lifespan: Extending A robust pipeline for functional MRI preprocessing to developmental neuroimaging. In bioRxiv (p. 2025.05.14.654069). 10.1101/2025.05.14.654069

Gordon, E. M., Laumann, T. O., Gilmore, A. W., Newbold, D. J., Greene, D. J., Berg, J. J., Ortega, M., Hoyt-Drazen, C., Gratton, C., Sun, H., Hampton, J. M., Coalson, R. S., Nguyen, A. L., McDermott, K. B., Shimony, J. S., Snyder, A. Z., Schlaggar, B. L., Petersen, S. E., Nelson, S. M., & Dosenbach, N. U. F. (2017). Precision Functional Mapping of Individual Human Brains. Neuron, 95(4), 791–807.e7.

Graham, A. M., Buss, C., Rasmussen, J. M., Rudolph, M. D., Demeter, D. V., Gilmore, J. H., Styner, M., Entringer, S., Wadhwa, P. D., & Fair, D. A. (2016). Implications of newborn amygdala connectivity for fear and cognitive development at 6-months-of-age. Developmental Cognitive Neuroscience, 18, 12–25.

Gratton, C., Kraus, B. T., Greene, D. J., Gordon, E. M., Laumann, T. O., Nelson, S. M., Dosenbach, N. U. F., & Petersen, S. E. (2020). Defining Individual-Specific Functional Neuroanatomy for Precision Psychiatry. Biological Psychiatry, 88(1), 28–39.

Gratton, C., Laumann, T. O., Nielsen, A. N., Greene, D. J., Gordon, E. M., Gilmore, A. W., Nelson, S. M., Coalson, R. S., Snyder, A. Z., Schlaggar, B. L., Dosenbach, N. U. F., & Petersen, S. E. (2018). Functional Brain Networks Are Dominated by Stable Group and Individual Factors, Not Cognitive or Daily Variation. Neuron, 98(2), 439–452.e5.

Hendrickson, T. J., Reiners, P., Moore, L. A., Lundquist, J. T., Fayzullobekova, B., Perrone, A. J., Lee, E. G., Moser, J., Day, T. K. M., Alexopoulos, D., Styner, M., Kardan, O., Chamberlain, T. A., Mummaneni, A., Caldas, H. A., Bower, B., Stoyell, S., Martin, T., Sung, S., … Feczko, E. (2024). BIBSNet: A deep learning Baby image brain segmentation network for MRI scans. In bioRxivorg (p. 2023.03.22.533696). 10.1101/2023.03.22.533696

Hendrikx, D., Smits, A., Lavanga, M., De Wel, O., Thewissen, L., Jansen, K., Caicedo, A., Van Huffel, S., & Naulaers, G. (2019). Measurement of neurovascular coupling in neonates. Frontiers in Physiology, 10, 65.

Hermosillo, R. J. M., Moore, L. A., Feczko, E., Miranda-Domínguez, Ó., Pines, A., Dworetsky, A., Conan, G., Mooney, M. A., Randolph, A., Graham, A., Adeyemo, B., Earl, E., Perrone, A., Carrasco, C. M., Uriarte-Lopez, J., Snider, K., Doyle, O., Cordova, M., Koirala, S., … Fair, D. A. (2024). A precision functional atlas of personalized network topography and probabilities. Nature Neuroscience, 1–14.

Hu, H., Cusack, R., & Naci, L. (2022). Typical and disrupted brain circuitry for conscious awareness in full-term and preterm infants. Brain Communications, 4(2), fcac071.

Hutchison, R. M., Culham, J. C., Flanagan, J. R., Everling, S., & Gallivan, J. P. (2015). Functional subdivisions of medial parieto-occipital cortex in humans and nonhuman primates using resting-state fMRI. NeuroImage, 116, 10–29.

Jiang, Y., Komatsu, M., Chen, Y., Xie, R., Zhang, K., Xia, Y., Gui, P., Liang, Z., & Wang, L. (2022). Constructing the hierarchy of predictive auditory sequences in the marmoset brain. eLife, 11, e74653.

Katus, L., Mason, L., Milosavljevic, B., McCann, S., Rozhko, M., Moore, S. E., Elwell, C. E., Lloyd-Fox, S., de Haan, M., & BRIGHT project team. (2020). ERP markers are associated with neurodevelopmental outcomes in 1-5 month old infants in rural Africa and the UK. NeuroImage, 210(116591), 116591.

Kim, H. (2014). Involvement of the dorsal and ventral attention networks in oddball stimulus processing: a meta-analysis. Human Brain Mapping, 35(5), 2265–2284.

King, G., Truzzi, A., & Cusack, R. (2023). The confound of head position in within-session connectome fingerprinting in infants. NeuroImage, 265, 119808.

Korom, M., Camacho, M. C., Filippi, C. A., Licandro, R., Moore, L. A., Dufford, A., Zöllei, L., Graham, A. M., Spann, M., Howell, B., FIT’NG, Shultz, S., & Scheinost, D. (2021). Dear reviewers: Responses to common reviewer critiques about infant neuroimaging studies. Developmental Cognitive Neuroscience, 53, 101055.

Kosakowski, H. L., Norman-Haignere, S., Mynick, A., Takahashi, A., Saxe, R., & Kanwisher, N. (2023). Preliminary evidence for selective cortical responses to music in one-month-old infants. Developmental Science, 26(5), e13387.

Labonte, A. K., Camacho, M. C., Moser, J., Koirala, S., Laumann, T. O., Marek, S., Fair, D., & Sylvester, C. M. (2024). Precision functional mapping to advance developmental psychiatry research. Biological Psychiatry Global Open Science, 100370, 100370.

Labonte, A. K., Moser, J., Camacho, M. C., Tu, J. C., Wheelock, M., Laumann, T. O., Gordon, E. M., Fair, D. A., & Sylvester, C. (2025). Precision functional mapping of the individual human brain near birth. In bioRxiv (p. 2025.07.07.663543). 10.1101/2025.07.07.663543

Laemle, L. K., Strominger, N. L., & Carpenter, D. O. (2006). Cross-modal innervation of primary visual cortex by auditory fibers in congenitally anophthalmic mice. Neuroscience Letters, 396(2), 108–112.

Laumann, T. O., Gordon, E. M., Adeyemo, B., Snyder, A. Z., Joo, S. J., Chen, M.-Y., Gilmore, A. W., McDermott, K. B., Nelson, S. M., Dosenbach, N. U. F., Schlaggar, B. L., Mumford, J. A., Poldrack, R. A., & Petersen, S. E. (2015). Functional System and Areal Organization of a Highly Sampled Individual Human Brain. Neuron, 87(3), 657–670.

Leo, A., Bernardi, G., Handjaras, G., Bonino, D., Ricciardi, E., & Pietrini, P. (2012). Increased BOLD variability in the parietal cortex and enhanced parieto-occipital connectivity during tactile perception in congenitally blind individuals. Neural Plasticity, 2012(1), 720278.

Liu, Y., Gao, J. H., Liu, H. L., & Fox, P. T. (2000). The temporal response of the brain after eating revealed by functional MRI. Nature, 405(6790), 1058–1062.

Lynch, C. J., Elbau, I. G., Ng, T., Ayaz, A., Zhu, S., Wolk, D., Manfredi, N., Johnson, M., Chang, M., Chou, J., Summerville, I., Ho, C., Lueckel, M., Bukhari, H., Buchanan, D., Victoria, L. W., Solomonov, N., Goldwaser, E., Moia, S., … Liston, C. (2024). Frontostriatal salience network expansion in individuals in depression. Nature, 1–10.

Marek, S., & Greene, D. J. (2021). Precision Functional Mapping of the Subcortex and Cerebellum. Current Opinion in Behavioral Sciences, 40, 12–18.

Mehler, J., Jusczyk, P., Lambertz, G., Halsted, N., Bertoncini, J., & Amiel-Tison, C. (1988). A precursor of language acquisition in young infants. Cognition, 29(2), 143–178.

Menz, M. M., Neumann, J., Müller, K., & Zysset, S. (2006). Variability of the BOLD response over time: an examination of within-session differences. NeuroImage, 32(3), 1185–1194.

Moeller, S., Pisharady, P. K., Ramanna, S., Lenglet, C., Wu, X., Dowdle, L., Yacoub, E., Uğurbil, K., & Akçakaya, M. (2021). NOise reduction with DIstribution Corrected (NORDIC) PCA in dMRI with complex-valued parameter-free locally low-rank processing. NeuroImage, 226, 117539.

Moeller, S., Yacoub, E., Olman, C. A., Auerbach, E., Strupp, J., Harel, N., & Uğurbil, K. (2010). Multiband multislice GE-EPI at 7 tesla, with 16-fold acceleration using partial parallel imaging with application to high spatial and temporal whole-brain fMRI. Magnetic Resonance in Medicine: Official Journal of the Society of Magnetic Resonance in Medicine / Society of Magnetic Resonance in Medicine, 63(5), 1144–1153.

Molloy, M. F., & Saygin, Z. M. (2022). Individual variability in functional organization of the neonatal brain. NeuroImage, 253, 119101.

Moore, L. A., Hermosillo, R. J. M., Feczko, E., Moser, J., Koirala, S., Allen, M. C., Buss, C., Conan, G., Juliano, A. C., Marr, M., Miranda-Dominguez, O., Mooney, M., Myers, M., Rasmussen, J., Rogers, C. E., Smyser, C. D., Snider, K., Sylvester, C., Thomas, E., … Graham, A. M. (2024). Towards personalized precision functional mapping in infancy. Imaging Neuroscience. 10.1162/imag_a_00165

Moser, J., Nelson, S. M., Koirala, S., Madison, T. J., Labonte, A. K., Carrasco, C. M., Feczko, E., Moore, L. A., Lundquist, J. T., Weldon, K. B., Grimsrud, G., Hufnagle, K., Ahmed, W., Myers, M. J., Adeyemo, B., Snyder, A. Z., Gordon, E. M., Dosenbach, N. U. F., Tervo-Clemmens, B., … Fair, D. A. (2024). Multi-echo acquisition and thermal denoising advances precision functional imaging. Imaging Neuroscience, 3, imag_a_00426.

Moser, J., Schleger, F., Weiss, M., Sippel, K., Dehaene-Lambertz, G., & Preissl, H. (2020). Magnetoencephalographic signatures of hierarchical rule learning in newborns. Developmental Cognitive Neuroscience, 46(100871), 100871.

Moser, J., Schleger, F., Weiss, M., Sippel, K., Semeia, L., & Preissl, H. (2021). Magnetoencephalographic signatures of conscious processing before birth. Developmental Cognitive Neuroscience, 49(100964), 100964.

Myers, M. J., Labonte, A. K., Gordon, E. M., Laumann, T. O., Tu, J. C., Wheelock, M. D., Nielsen, A. N., Schwarzlose, R. F., Camacho, M. C., Alexopoulos, D., Warner, B. B., Raghuraman, N., Luby, J. L., Barch, D. M., Fair, D. A., Petersen, S. E., Rogers, C. E., Smyser, C. D., & Sylvester, C. M. (2024). Functional parcellation of the neonatal cortical surface. Cerebral Cortex, 34(2). 10.1093/cercor/bhae047

Nilearn contributors, Chamma, A., Frau-Pascual, A., Rothberg, A., Abadie, A., Abraham, A., Gramfort, A., Savio, A., Cionca, A., Sayal, A., Thual, A., Kodibagkar, A., Kanaan, A., Pinho, L., Joshi, A., Idrobo, A. H., Kieslinger, A.-S., Kumari, A., Rokem, A., … Nájera, Ó. (2025). nilearn. Zenodo. 10.5281/ZENODO.14697221

Prince, J. S., Charest, I., Kurzawski, J. W., Pyles, J. A., Tarr, M. J., & Kay, K. N. (2022). Improving the accuracy of single-trial fMRI response estimates using GLMsingle. eLife, 11. 10.7554/eLife.77599

Rieck, J. R., DeSouza, B., Baracchini, G., & Grady, C. L. (2022). Reduced modulation of BOLD variability as a function of cognitive load in healthy aging. Neurobiology of Aging, 112, 215–230.

Rogers, C. E., Sylvester, C. M., Mintz, C., Kenley, J. K., Shimony, J. S., Barch, D. M., & Smyser, C. D. (2017). Neonatal Amygdala Functional Connectivity at Rest in Healthy and Preterm Infants and Early Internalizing Symptoms. Journal of the American Academy of Child and Adolescent Psychiatry, 56(2), 157–166.

Schleger, F., Landerl, K., Muenssinger, J., Draganova, R., Reinl, M., Kiefer-Schmidt, I., Weiss, M., Wacker-Gußmann, A., Huotilainen, M., & Preissl, H. (2014). Magnetoencephalographic signatures of numerosity discrimination in fetuses and neonates. Developmental Neuropsychology, 39(4), 316–329.

Siegel, J. S., Power, J. D., Dubis, J. W., Vogel, A. C., Church, J. A., Schlaggar, B. L., & Petersen, S. E. (2014). Statistical improvements in functional magnetic resonance imaging analyses produced by censoring high-motion data points. Human Brain Mapping, 35(5), 1981–1996.

Spann, M. N., Monk, C., Scheinost, D., & Peterson, B. S. (2018). Maternal Immune Activation During the Third Trimester Is Associated with Neonatal Functional Connectivity of the Salience Network and Fetal to Toddler Behavior. The Journal of Neuroscience: The Official Journal of the Society for Neuroscience, 38(11), 2877–2886.

Sylvester, C. M., Kaplan, S., Myers, M. J., Gordon, E. M., Schwarzlose, R. F., Alexopoulos, D., Nielsen, A. N., Kenley, J. K., Meyer, D., Yu, Q., Graham, A. M., Fair, D. A., Warner, B. B., Barch, D. M., Rogers, C. E., Luby, J. L., Petersen, S. E., & Smyser, C. D. (2022). Network-specific selectivity of functional connections in the neonatal brain. Cerebral Cortex. 10.1093/cercor/bhac202

Sylvester, C. M., Myers, M. J., Perino, M. T., Kaplan, S., Kenley, J. K., Smyser, T. A., Warner, B., Barch, D. M., Pine, D. S., Luby, J. L., Rogers, C. E., & Smyser, C. D. (2021). Neonatal Brain Response to Deviant Auditory Stimuli and Relation to Maternal Trait Anxiety. The American Journal of Psychiatry, 178(8), 771–778.

Sylvester, C. M., Smyser, C. D., Smyser, T., Kenley, J., Ackerman, J. J., Jr, Shimony, J. S., Petersen, S. E., & Rogers, C. E. (2018). Cortical Functional Connectivity Evident After Birth and Behavioral Inhibition at Age 2. The American Journal of Psychiatry, 175(2), 180–187.

Thoman, E. B., & Tynan, W. D. (1979). Sleep states and wakefulness in human infants: profiles from motility monitoring. Physiology & Behavior, 23(3), 519–525.

Thomason, M. E., Grove, L. E., Lozon, T. A., Jr, Vila, A. M., Ye, Y., Nye, M. J., Manning, J. H., Pappas, A., Hernandez-Andrade, E., Yeo, L., Mody, S., Berman, S., Hassan, S. S., & Romero, R. (2015). Age-related increases in long-range connectivity in fetal functional neural connectivity networks in utero. Developmental Cognitive Neuroscience, 11, 96–104.

Uhrig, L., Dehaene, S., & Jarraya, B. (2014). A hierarchy of responses to auditory regularities in the macaque brain. The Journal of Neuroscience: The Official Journal of the Society for Neuroscience, 34(4), 1127–1132.

Vizioli, L., Moeller, S., Dowdle, L., Akçakaya, M., De Martino, F., Yacoub, E., & Uğurbil, K. (2021). Lowering the thermal noise barrier in functional brain mapping with magnetic resonance imaging. Nature Communications, 12(1), 5181.

von Melchner, L., Pallas, S. L., & Sur, M. (2000). Visual behaviour mediated by retinal projections directed to the auditory pathway. Nature, 404(6780), 871–876.

Wacongne, C., Labyt, E., van Wassenhove, V., Bekinschtein, T., Naccache, L., & Dehaene, S. (2011). Evidence for a hierarchy of predictions and prediction errors in human cortex. Proceedings of the National Academy of Sciences of the United States of America, 108(51), 20754–20759.

Weber, P., Depoorter, A., Hetzel, P., & Lemola, S. (2016). Habituation as parameter for prediction of mental development in healthy preterm infants: An electrophysiological pilot study: An electrophysiological pilot study. Journal of Child Neurology, 31(14), 1591–1597.

Yates, T. S., Fel, J., Choi, D., Trach, J. E., Behm, L., Ellis, C. T., & Turk-Browne, N. B. (2025). Hippocampal encoding of memories in human infants. Science (New York, N.Y.), 387(6740), 1316–1320.

Zhang, Y., Deng, Q., Wang, J., Wang, H., Li, Q., Zhu, B., Ji, C., Xu, X., & Johnston, L. (2022). The impact of breast milk feeding on early brain development in preterm infants in China: An observational study. PloS One, 17(11), e0272125.

