## Supplementary Material for "Precision Functional Neuroimaging Reveals Individually Specific Auditory Responses in Infants"

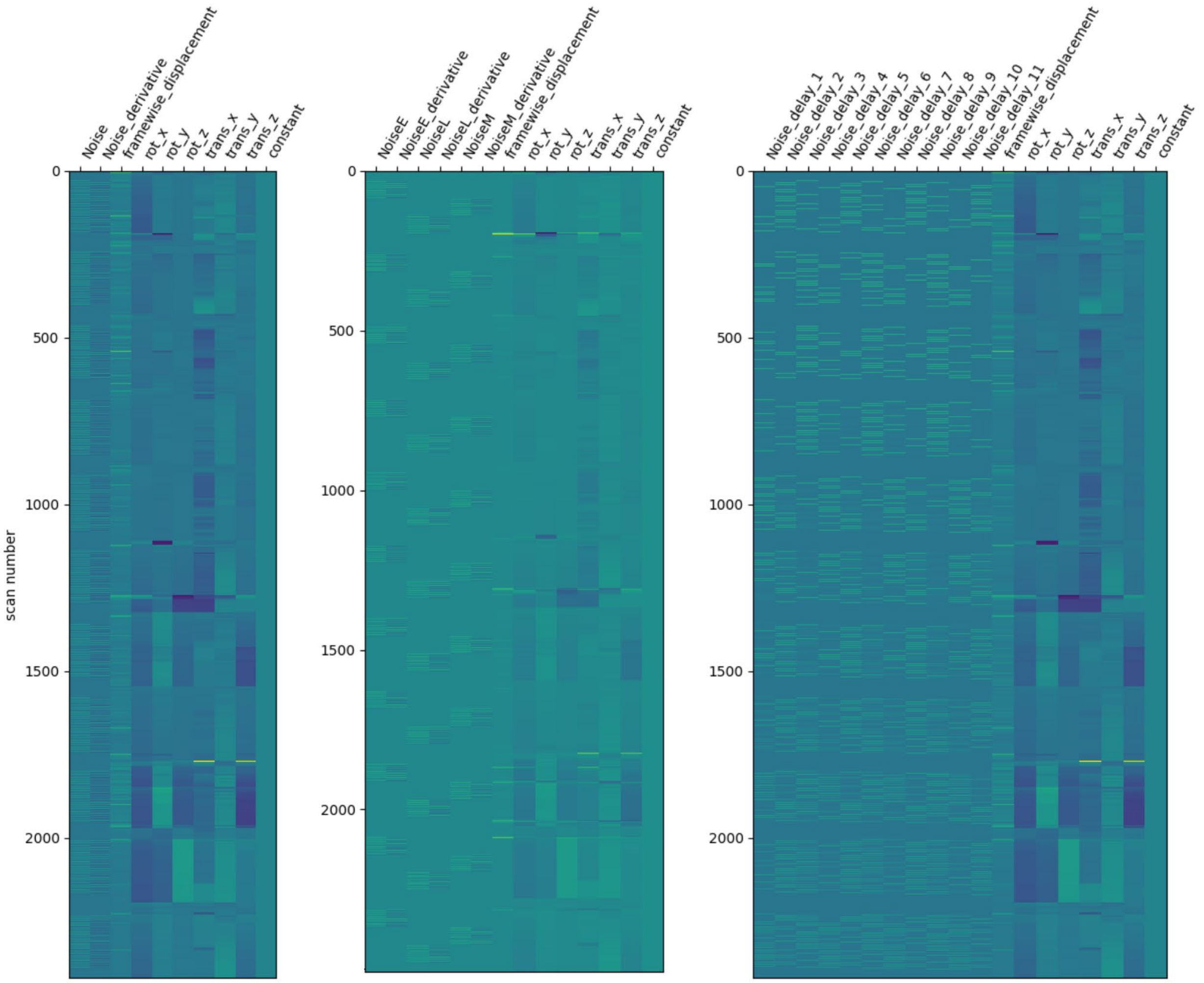


*Figure S1: Example design matrices for standard model, model with early, middle and late oddball coded and Finite Impulse Response model. Example participant PB0011.*

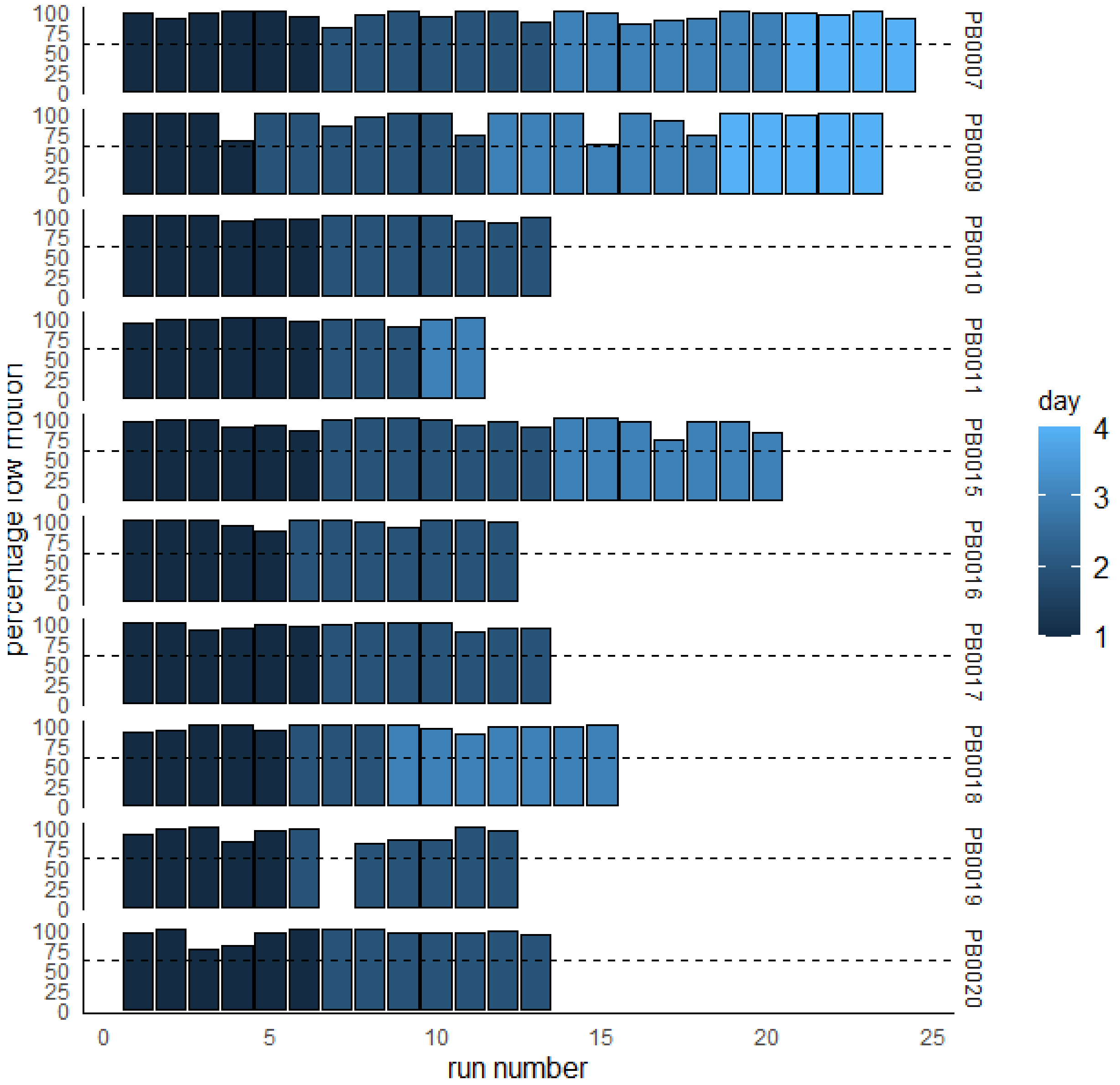


*Figure S2: Data overview showing all runs of each participant colored by acquisition day using task FD threshold (<0.9 mm). Dashed line represents 60% inclusion criteria.*

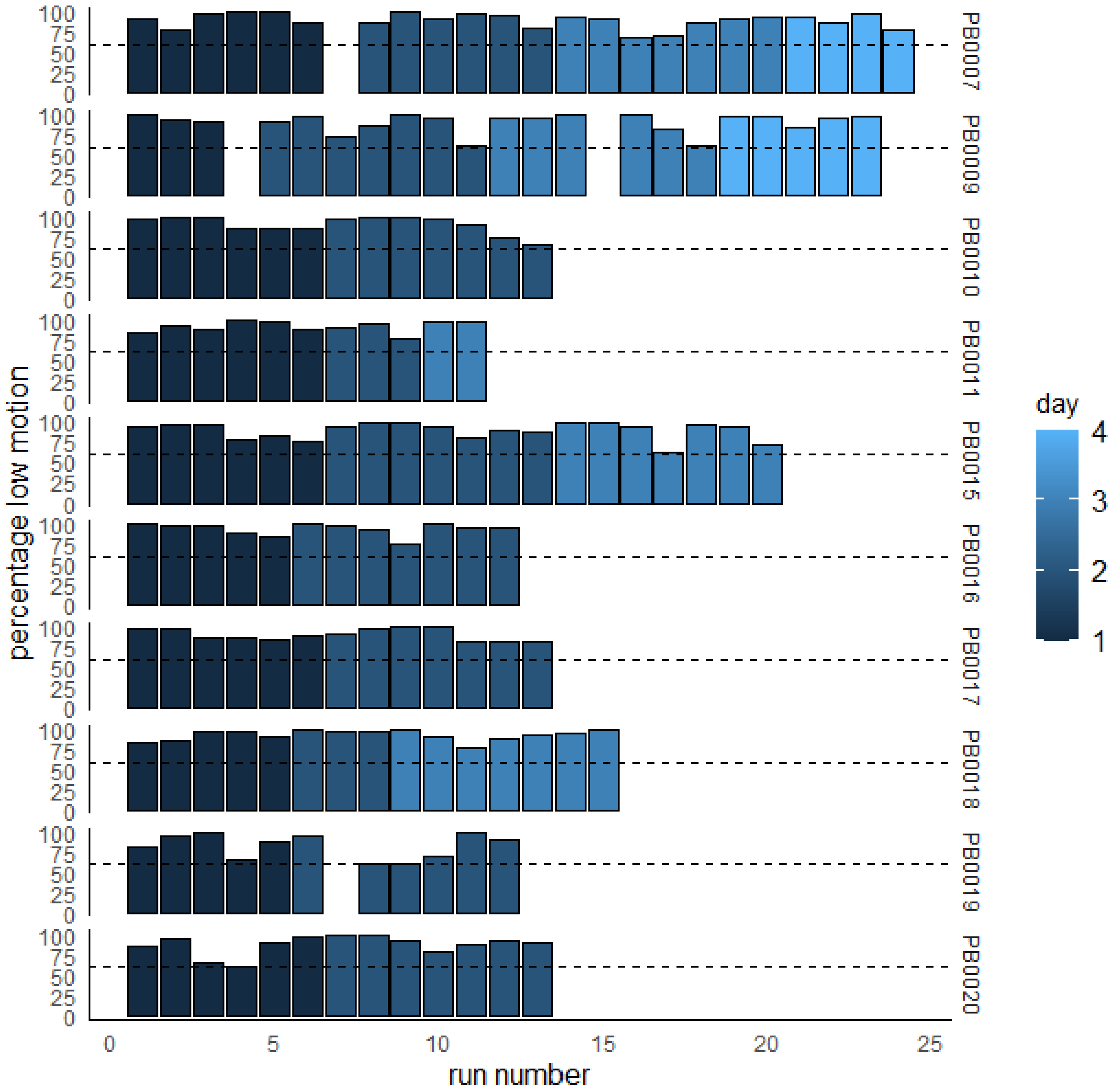


*Figure S3: Data overview showing all runs of each participant colored by acquisition day using connectivity analysis FD threshold (<0.3 mm). Dashed line represents 60% inclusion criteria.*

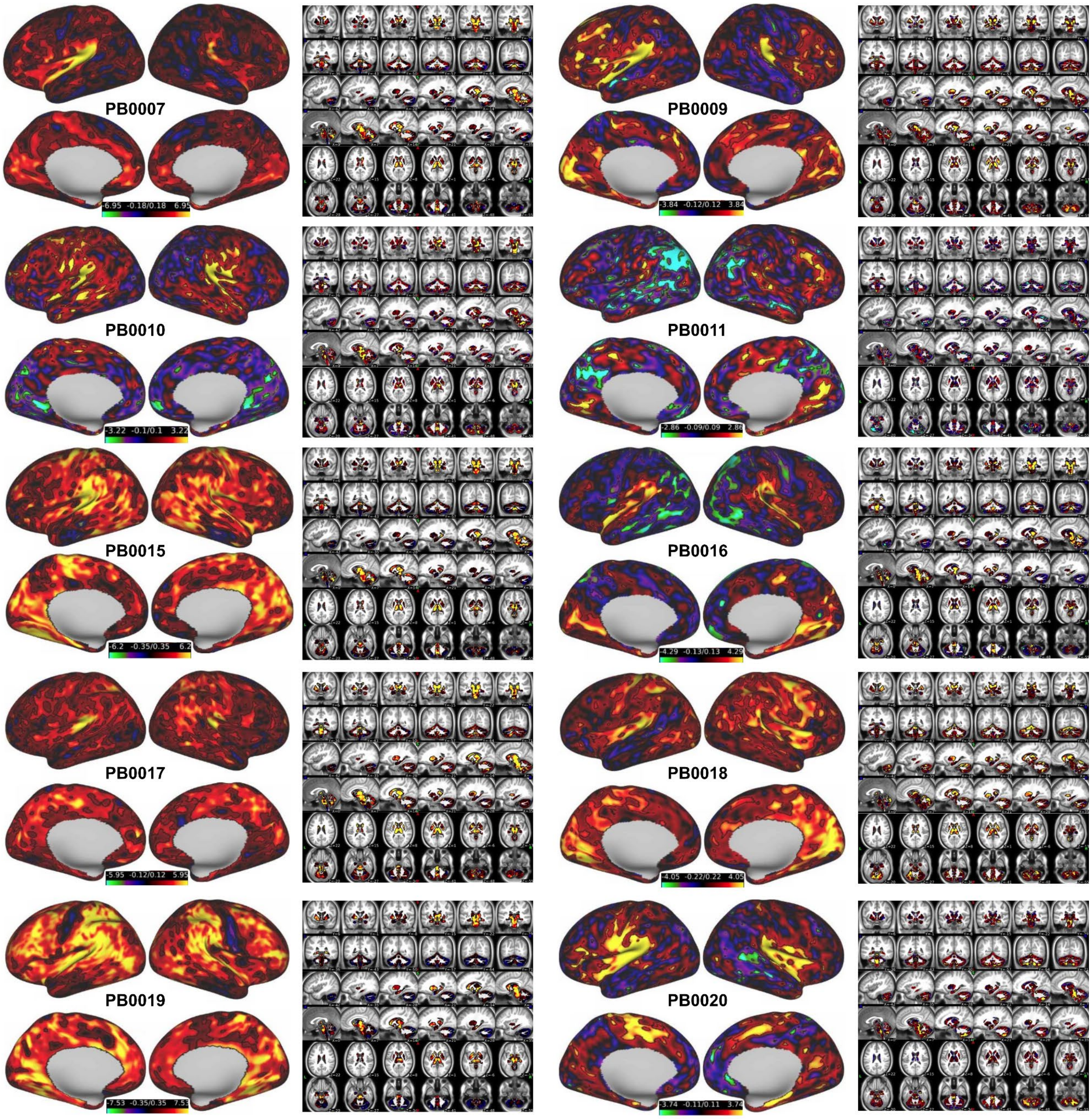


*Figure S4: Beta maps for each individual participant. Scales are adjusted to 5%-95% of individual beta values. Black outlines outline areas with significant beta values (p<0.01).*

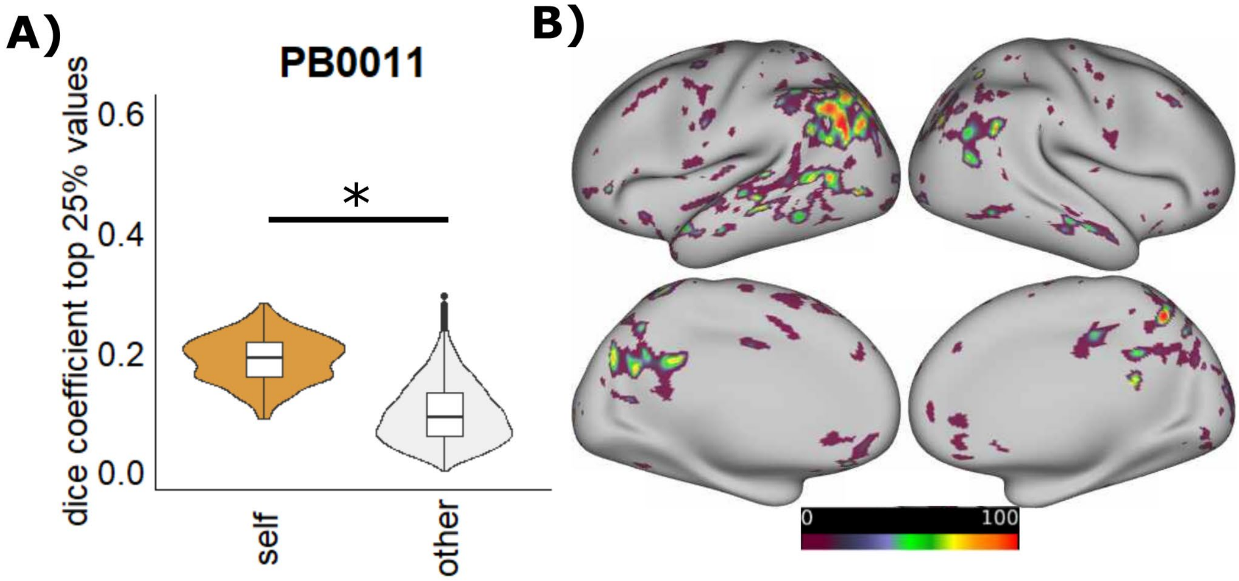


*Figure S5: A) Dice overlap of top 25% of negative beta values between split halves in 462 unique combinations of runs for PB0011. Comparisons are between split halves (colored) and the first half of every other subject (light gray). B) Overlap of split halves across permutations thresholded at the 25th percentile of negative betas. Scale is in percent.*

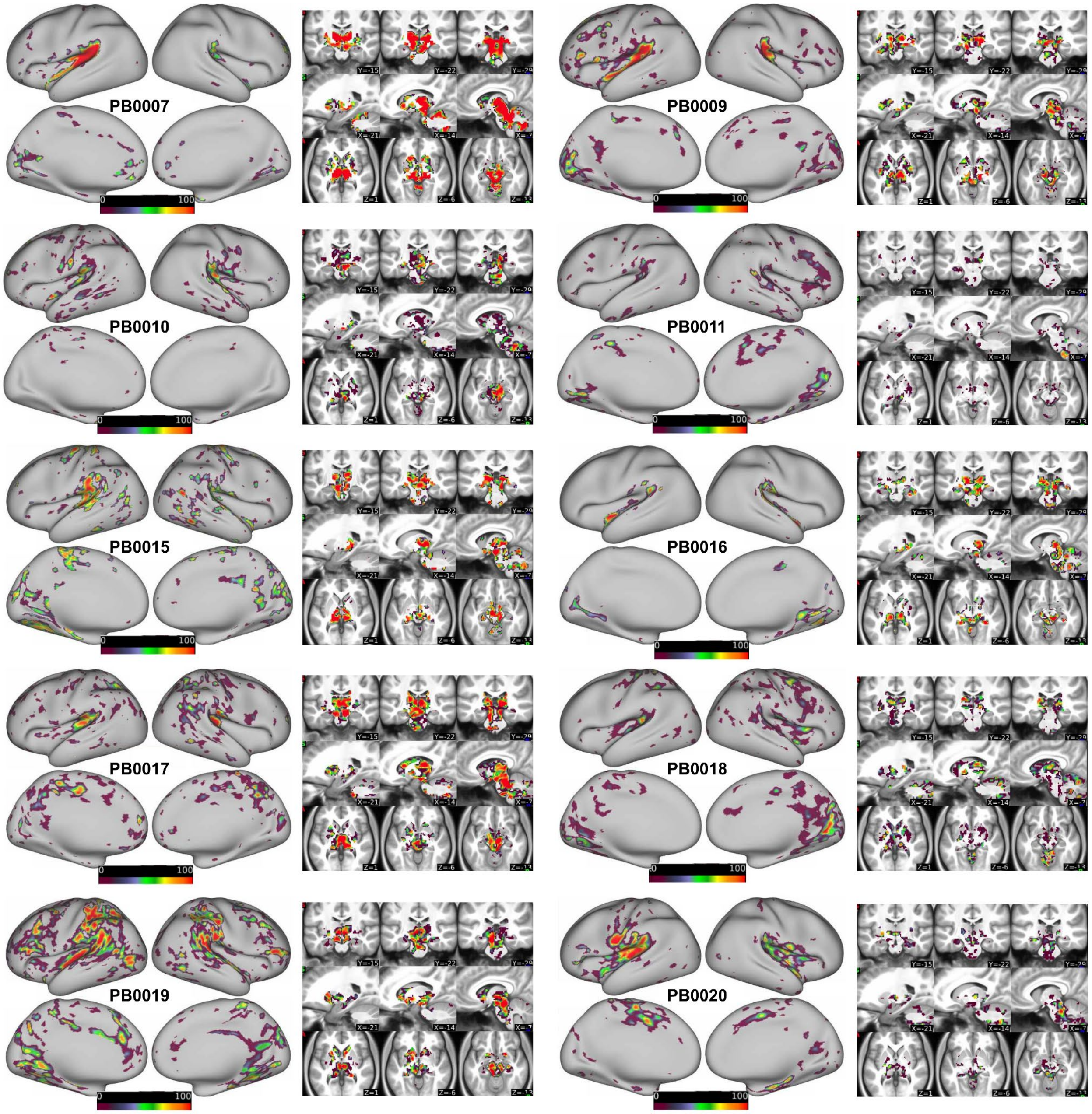


*Figure S6: Overlap of split halves across permutations thresholded at the 75th percentile. Scale is in percent. Maps represent areas that overlap between split-halves and across permutations.*

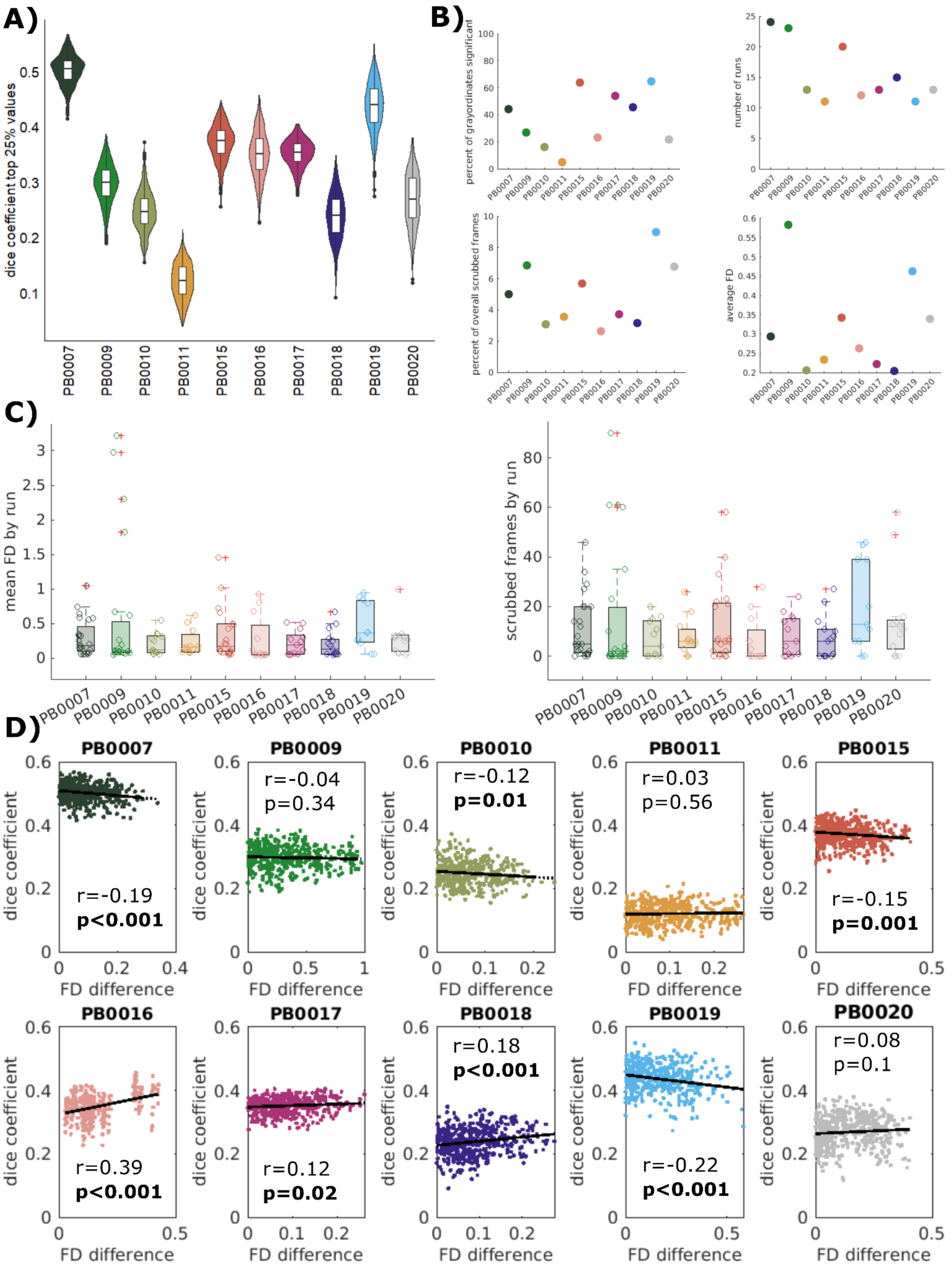
A)
dice coefficient top 25% values
0.1
0.2
0.3
0.4
0.5
PB0007
PB0009
PB0010
PB0011
PB0015
PB0016
PB0017
PB0018
PB0019
PB0020
B)
percent of grayordinates significant
number of runs
percent of overall scrubbed frames
average FD
C)
mean FD by run
scrubbed frames by run
D)
PB0007
r=-0.19
p<0.001
PB0009
r=-0.04
p=0.34
PB0010
r=-0.12
p=0.01
PB0011
r=0.03
p=0.56
PB0015
r=-0.15
p=0.001
PB0016
r=0.39
p<0.001
PB0017
r=0.12
p=0.02
PB0018
r=0.18
p<0.001
PB0019
r=-0.22
p<0.001
PB0020
r=0.08
p=0.1
dice coefficient
FD difference

*Figure S7: A) Overlap between split halves across permutations (summary of panels in Figure 2). B) Factors that could potentially contribute to differences between participants such as the number of runs, the average FD and the number of scrubbed frames from the data. There is no apparent relationship between these variables and the split-half stability. However, the percentage of significant vertices mostly lines up with the subject's stability metric C) More fine grained display of average FD and number of scrubbed frames by run D) Relationship between overlap between split halves and mean FD of each of the halves quantified by the FD difference between halves (absolute value). For some participants run combinations in which runs with higher motion are all grouped in one half of the split-half (high FD difference) tend to show lower dice coefficients (negative r) while others show the opposite effect or no relationship.*

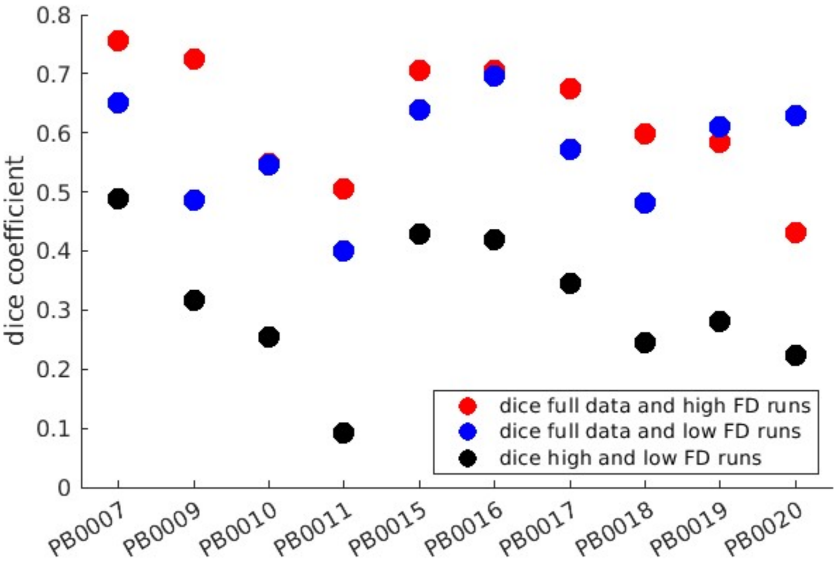


*Figure S8: Overlap of 75% highest beta values from all data or half of a participant's data, taking the run combination with the highest motion or lowest motion. High motion runs do not significantly contribute more to the average (t-test between dice with high and low motion runs t(9)=1.41, p=0.19). Dice values are higher than in previous figures as this analysis includes overlapping runs.*

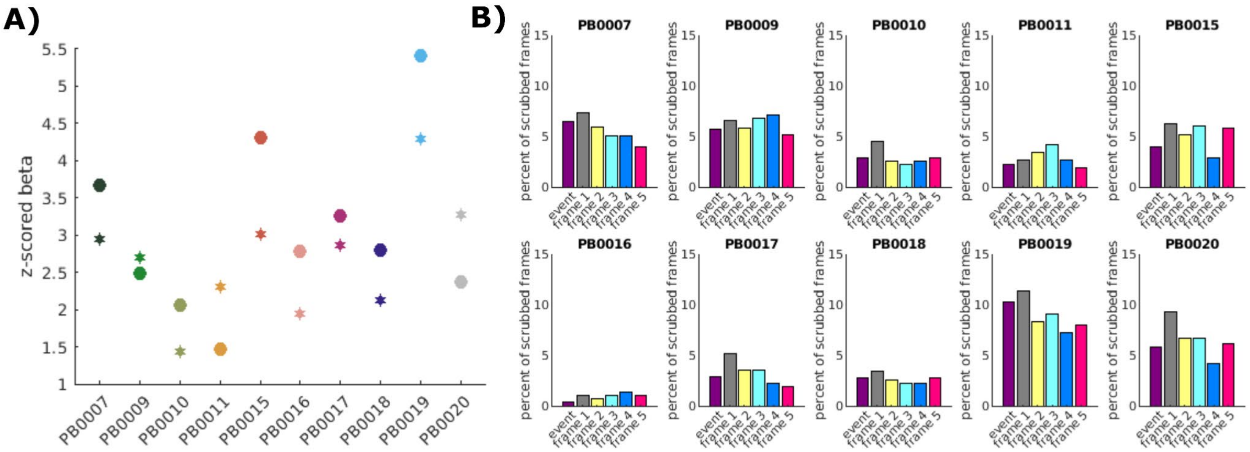


*FigureS9: A) Comparison of 75th percentile of beta values from model with assumed response (circles) to model with Finite Impulse Response (FIR; stars). For the FIR model, maximal beta values were extracted from the 11 modeled frames (~ 20 seconds) after oddball onset. Maximal values were selected independent of timing and could be in any of the 11 frames. From these maximal values, the 75th percentile is plotted. Despite values being closer together, variability in beta values remains with FIR model and rank between participants is similar. B) percentage of frames that were removed after onset of the oddball event for each participant. Fraction of removed frames is low for all participants and patterns not systematically related to beta magnitudes.*

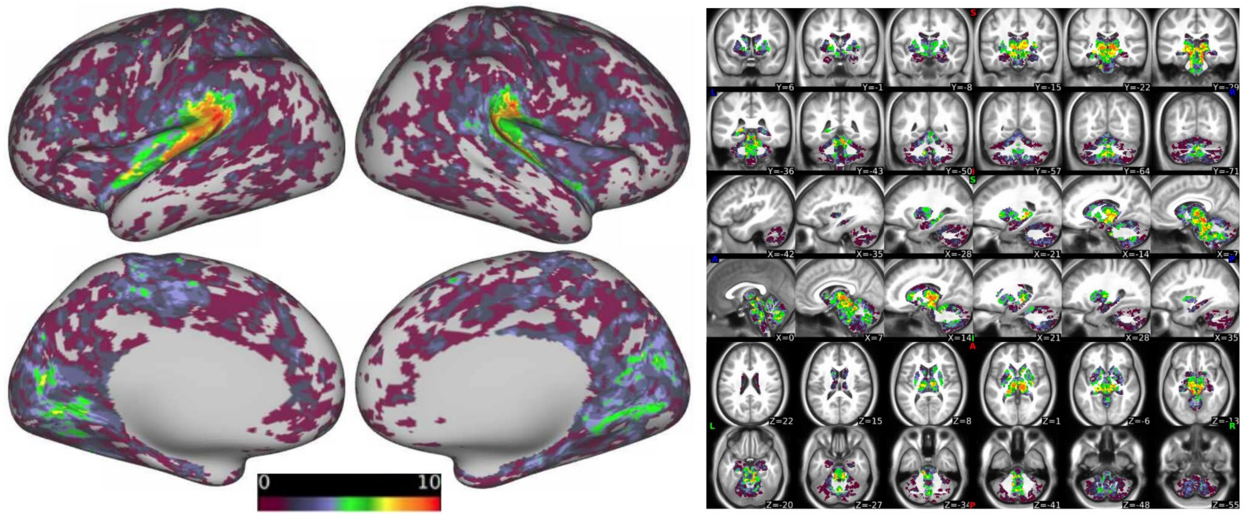


*FigureS10: Overlap of areas with highest 25% beta values across individuals. Scale 0-10 is the participant count. Overlap shows similar pattern as in Figure 3.*

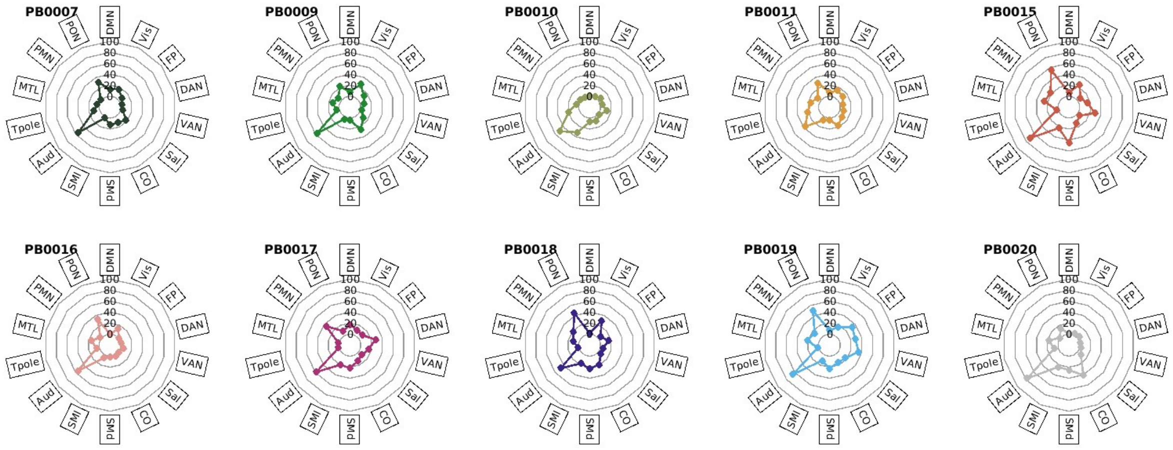


*Figure S11: Individual specific depiction of Figure 5. Percent of a network containing high beta values (top 25%) for each participant. DMN: default mode network; Vis: visual network; FP: fronto-parietal network; DAN: dorsal attention network; VAN: ventral attention network; Sal: salience network; CO: cingulo-opercular network; SMd: somato-motor dorsal; SMl: somato-motor lateral; Aud: auditory network; Tpole: temporal pole; MTL: medial temporal lobe; PMN: parietal memory network; PON: parietal-occipital network.*

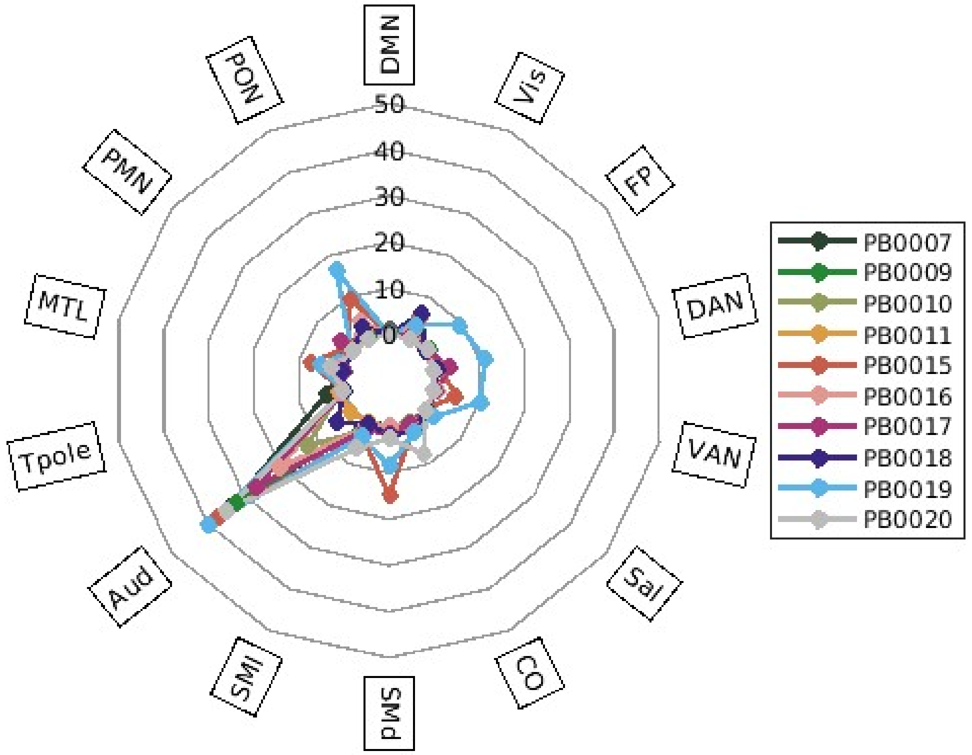


*Figure S12: Figure 5 restricted to areas that overlap between split halves in more than 50% of permutations.*

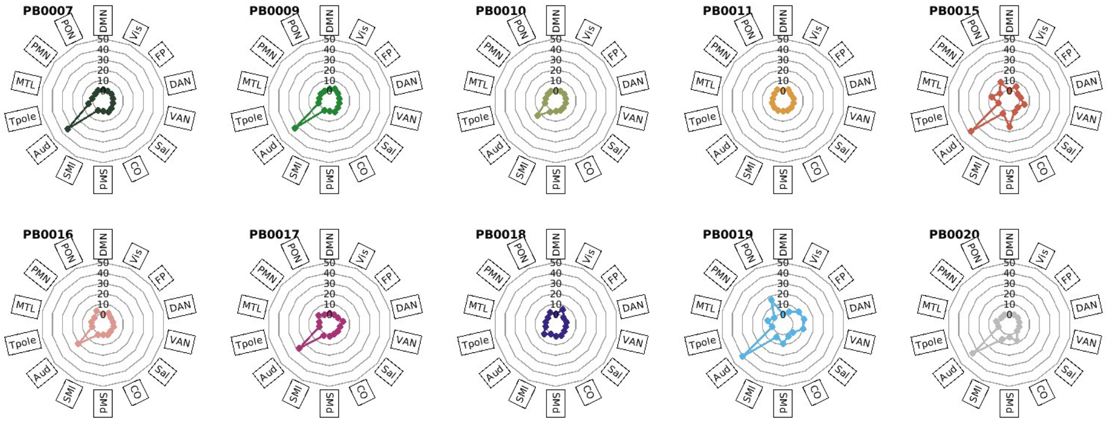


*Figure S13: Figure S11 restricted to areas that overlap between split halves in more than 50% of permutations.*

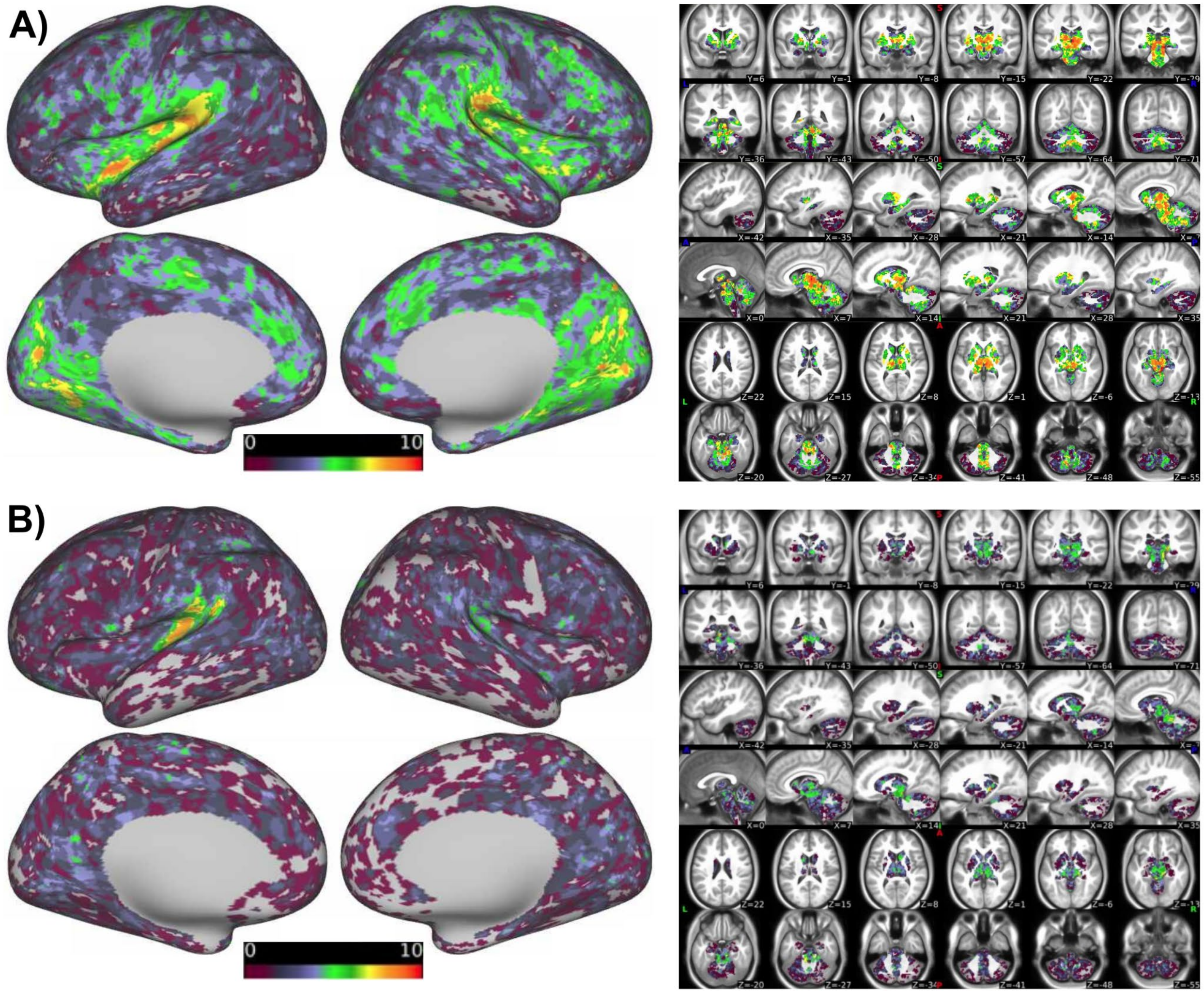


*Figure S14: Overlap of areas with significant activation (p<0.01) across individuals. Scale 0-10 is the participant count. A) early (first 8) oddballs within a run only B) late (last 8) oddballs within a run only.*
